# A neural signature of choice under sensory conflict in *Drosophila*

**DOI:** 10.1101/2020.08.14.251553

**Authors:** Preeti Sareen, Li Yan McCurdy, Michael N. Nitabach

## Abstract

Feeding decisions are fundamental to survival, and decision making is often disrupted in disease, yet the neuronal and molecular mechanisms of adaptive decision making are not well understood. Here we show that neural activity in a small population of neurons projecting to the fan-shaped body higher-order central brain region of *Drosophila* represents final food choice during sensory conflict. We found that hungry flies made tradeoffs between appetitive and aversive values of food in a decision making task to choose bittersweet food with high sucrose concentration, but adulterated with bitter quinine, over sweet-only food with less sucrose. Using cell-specific optogenetics and receptor RNAi knockdown during the decision task, we identified an upstream neuropeptidergic and dopaminergic network that relays internal state and other decision-relevant information, such as valence and previous experience, to a specific subset of fan-shaped body neurons. Importantly, calcium imaging revealed that these neurons were strongly inhibited by the taste of the rejected food choice, suggesting that they encode final behavioral food choice. Our findings reveal that fan-shaped body taste responses to food choices are determined not only by taste quality, but also by previous experience (including choice outcome) and hunger state, which are integrated in the fan-shaped body to encode the decision before relay to downstream motor circuits for behavioral implementation. Our results uncover a novel role for the fan-shaped body in choice encoding, and reveal a neural substrate for sensory and internal state integration for decision making in a genetically tractable model organism to enable mechanistic dissection at circuit, cellular, and molecular levels.

## Introduction

Animals integrate food-related sensory information from their external environment with their internal state in order to make adaptive decisions. Often food-related sensory information is conflicting in valence. For example, *Drosophila* flies forage on decomposing fruits and must balance obtaining essential nutrition with avoiding toxins, pathogens, etc. As flies forage, sweet and bitter taste receptors on their legs and wings signal the presence of sweet nutritive food and bitter potential toxins (Scott, 2018). Flies must adaptively weigh and integrate this conflicting information before consumption to enhance survival fitness. While some studies have shown integration of sweet and bitter tastes at the sensory neuron level (Chu et al., 2014; Jeong et al., 2013; Meunier et al., 2003), very little is known about how taste and other external sensory cues are integrated with internal state of the animal to form a feeding decision in the higher brain. We investigated how value-based decisions are made in the central brain of a hungry fly, using an experimental paradigm in which freely foraging flies sample and choose between different sweet and bittersweet foods (Fig. 1A). We quantified food choice and manipulated subsets of neurons while flies engaged in this decision task with conflicting taste information (Fig. 1A).

**Figure 1.**
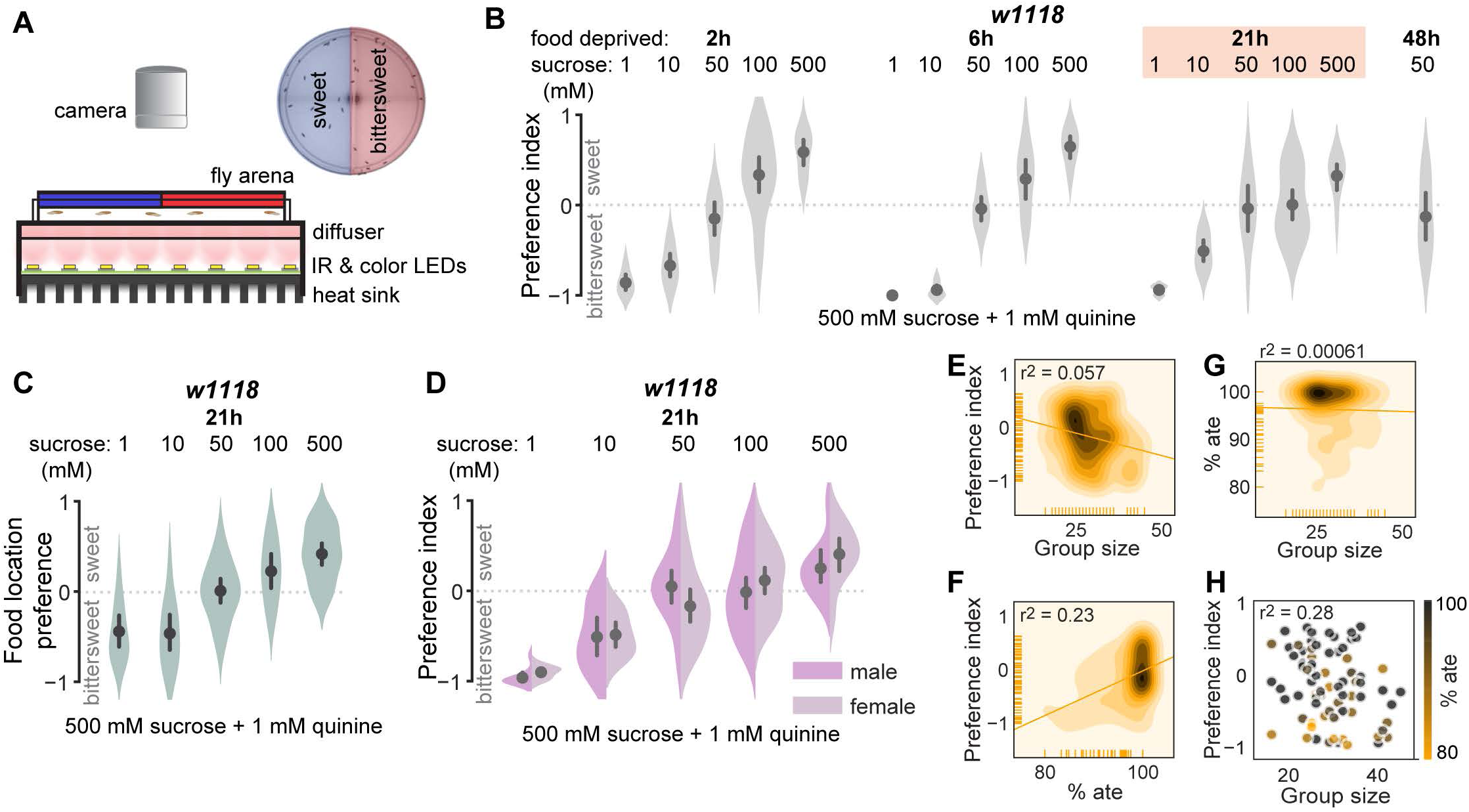
Hungry flies make trade-offs between the appetitive and aversive value of food. **A**, Schematic of the two-choice decision making assay. Sweet and bittersweet foods are prepared in dissolved agarose, mixed with food dyes (e.g., sweet with blue and bittersweet with red), and gelled in a circular arena. Dye colors are counterbalanced within each food tradeoff condition. Flies are introduced into the dark food arena to walk, sample, and consume freely for 5 min, while they are video recorded with infrared (IR) backlighting. Flies are then anaesthetized and abdomen color is recorded under a dissecting microscope, indicating ingested food choice. Preference index is calculated as (no. of sweet color flies + 0.5 purple flies) - (no. of bittersweet color flies + 0.5 purple flies)/total no. of flies that ate, where positive index means flies preferred sweet and negative index means flies preferred bittersweet. **B**, Preference index of wild-type (*w1118*) flies deprived of food for increasing durations indicate the tradeoff between the sweet and bittersweet food and reveal equal preference when the choice is between 50 mM sucrose-only and 500 mM sucrose + 1mM quinine. **C**, Position preference index (i.e., sweet or bittersweet sector preference as defined by the location of the flies at the end of the assay) matches ingested food preference, with equal preference at 50 mM sucrose-only vs. 500 mM sucrose + 1 mM quinine. **D,** Preferences of male and female flies within each group were indistinguishable for all conditions tested. There was no significant correlation between preference index and group size per trial (**E**), preference index and % of flies that ate per trial (**F**), or % of flies that ate per trial and group size (**G**). **H,** Neither group size nor % flies that ate significantly predicted preference index in a multiple regression model, indicating no interaction between these measures. **B-D**, Plots show mean ± 95% CI, violins depict kernel density estimation of the underlying data distribution with the width of each violin scaled by the number of observations at that y-value. Each violin summarizes 10 ≤ trials ≤ 30 with mode = 10. **E-H**, Heatmaps depict bivariate distribution visualized using a kernel density estimation procedure; darker regions indicate higher data density. r^2^ is the square of Pearson’s coefficient. See Table S1 for sample size and statistics.

## Results

### Hungry flies make tradeoffs when faced with conflicting sensory information

We quantified the choices of wild-type flies deprived of food for different durations between a range of increasing concentration of sweet (sucrose-only) food option and a constant bittersweet (sucrose + quinine) option, with each option including either a red or blue food dye. When choosing between a low sucrose concentration sweet option and a high sucrose concentration bittersweet option, flies prefer higher sucrose bittersweet (Fig. 1B), as quantified by color of food ingested. As sucrose concentration of the sweet choice increased, flies increasingly preferred it over bittersweet. This dose-dependent change in preference suggests that when the sweet option reaches sufficient sucrose, the caloric advantage in choosing a less palatable bittersweet food is outweighed by the danger-avoidance advantage of the sweet option (Fig. 1B). In the absence of bitter, flies always chose the sweeter option (Fig. S1A), suggesting that there was no saturation of sweet taste sensation for behavioral discrimination even at the highest sucrose concentration tested (500 mM sucrose). Flies equally preferred the sweet and bittersweet options at 10-fold sucrose concentration ratio (Fig. 1B, 50 mM vs. 500 mM sucrose + 1 mM quinine). This equal-preference point was identical at all of the tested food deprivation durations (Fig. 1B), suggesting an external taste sensation and internal hunger state equilibrium at this concentration ratio. The equal-preference point depends on the sucrose concentration ratio between the two options and not absolute concentration (Fig. S1B). These results indicate that hungry flies tradeoff the appetitive (sweet) and aversive (bitter) values of food in making feeding decisions, and are consistent with known effects of bitter compounds on feeding, in which increasing bitterness of food decreases feeding(Meunier et al., 2003).

To further characterize fly behavior in the decision task and determine whether sensory cues other than taste could affect food choice, we also determined the spatial location of each fly at the end of the decision task. As expected, position preference mirrored ingested food preference (Fig. 1C). We also addressed whether size or sex ratio of groups of flies tested together influenced the decision, since social interactions driven by group size (Ramdya et al., 2017; Rooke et al., 2020) and sex ratio can affect fly behavior. There was no effect on ingested food preference of male-to-female ratio within groups of flies tested together (Fig. 1D). At the equal-preference condition (21h deprivation, 50 mM sucrose vs. 500 mM sucrose + 1 mM quinine), neither food preference and group size (Fig. 1E), food preference and percent of flies that ate (Fig. 1F), nor percent of flies that ate and group size (Fig. 1G) were correlated. There was also no significant prediction of preference index by group size or percent of flies that ate in a multiple regression model (Fig. 1H, Table S1), indicating no interaction between these variables in the decision task. These results suggest that taste is the most salient sensory cue affecting food choice in the decision assay.

### A decision making neuronal ensemble converges on the fan-shaped body

During foraging, animals estimate values of internal state and external sensory parameters, such as hunger level, valence of available foods, location of food, etc. These value estimates are then integrated to form a decision. While neural substrates integrating internal state and external stimuli in the *Drosophila* brain remain unclear, there are hints in the literature of cell-populations and brain regions that could be involved. Various neuromodulators and the neurons that secrete them regulate hunger dependent food intake (Cannell et al., 2016; Hentze et al., 2015; Lin et al., 2019; Nassel and Zandawala, 2019; Wu et al., 2003; Yang et al., 2018; Zandawala et al., 2018), reward (Huetteroth et al., 2015; Liu et al., 2012a; Lyutova et al., 2019; Yamagata et al., 2015) or punishment (Riemensperger et al., 2005), and memory (Huetteroth et al., 2015; Masek et al., 2015; Yamagata et al., 2015). The mushroom body is an insect central brain region involved in gustatory learning and memory (Kirkhart and Scott, 2015; Masek and Scott, 2010) and valence encoding (Aso et al., 2014), and is thought to be a major center controlling higher-order behaviors, including associative learning (Cohn et al., 2015; Lewis et al., 2015; Tsao et al., 2018). The insect central complex is an evolutionarily conserved central brain region whose ellipsoid body and protocerebral bridge sub-regions have been implicated in navigation (Giraldo et al., 2018; Green et al., 2019; Honkanen et al., 2019; Kim et al., 2017b; Mathejczyk and Wernet, 2019; Seelig and Jayaraman, 2015; Sun et al., 2017; Turner-Evans et al., 2017) and sleep (Guo et al., 2018; Liu et al., 2019; Liu et al., 2016). The central complex fan-shaped body, a laminar sub-region, has been implicated in sleep (Donlea et al., 2014; Donlea et al., 2018; Liu et al., 2012b; Pimentel et al., 2016) and ethanol preference (Azanchi et al., 2013; Scaplen et al., 2020). Various neuromodulators (Kahsai and Winther, 2011), their receptors (Al-Anzi et al., 2010; Cavey et al., 2016; Donlea et al., 2018), and dopaminergic inputs (Kim et al., 2020; Qian et al., 2017) co-localize in the layers of fan-shaped body. We hypothesized that value estimates of internal state and external environment encoded by neuromodulatory neurons will be required inputs to higher-order brain regions for integration and generation of decisions. To test this hypothesis, we genetically targeted acute optogenetic activation and inhibition to specific neural subsets in neuromodulatory and higher-order brain centers using the GAL4-UAS binary expression system (Brand and Perrimon, 1993), while flies actively sampled and consumed food at the equal-preference condition (Fig. 1A, 1B, 21h food deprivation, 50 mM sucrose vs 500mM sucrose + 1mM quinine). For optogenetic activation, we used red-light-sensitive CsChrimson channelrhodopsin (Klapoetke et al., 2014), and for optogenetic inhibition we used green-light-sensitive anion-conducting channelrhodopsin GtACR1 (Mohammad et al., 2017). This optogenetic interrogation of modulatory neurons and higher order brain regions revealed neuropeptidergic neurons (Leucokinin, Allatostatin A, NPF, DH44), subsets of dopaminergic neurons (PPL1-γ2α’1, PPL1-α3, PAM-α1), and a narrow subset of fan-shaped body layer 6 neurons (FBl6) whose activation or inhibition significantly shifted food choice in the equal-preference condition (Fig. 2A).

**Figure 2.**
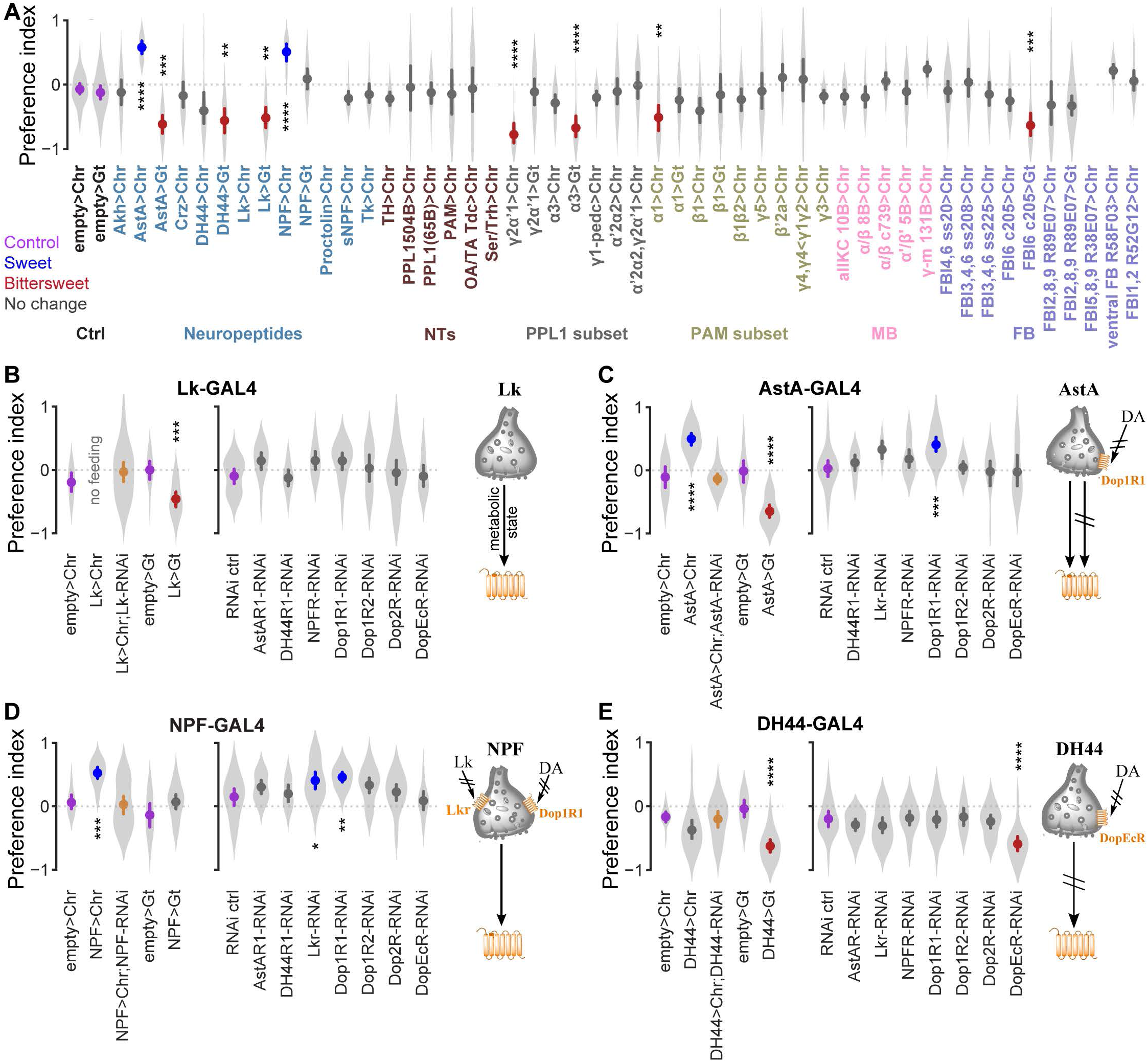
A decision making neuronal ensemble revealed by combined optogenetic and chemoconnectomic strategy. **A**, Cell-specific optogenetic activation and inhibition screen was performed at the equal preference condition (50 mM sucrose vs. 500 mM sucrose + 1 mM quinine) using flies deprived of food for 21 h. Neuronal subsets were genetically targeted using the GAL4-UAS binary expression system. Red-light-sensitive CsChrimson (Chr) was used for optogenetic activation and green-light-sensitive GtACR1 (Gt) for optogenetic silencing. Optogenetic manipulation of AstA, DH44, Lk, and NPF neuropeptides; PPL1 γ2α’1, PPL1 α3, and PAM α1 dopaminergic subsets; and fan-shaped body layer 6 neurons (FBl6) significantly shifted preference away from equal preference compared to respective empty>Chr and empty>Gt controls. Purple mean point = control, blue mean point = significant sweet preference, red mean point = significant bittersweet preference, yellow mean point = simultaneous optogenetic activation and RNAi. **B** (left), Optogenetic activation of Leucokinin (Lk) neurons suppressed feeding in food deprived flies, while optogenetic inhibition shifted the preference towards bittersweet. Simultaneous Lk RNAi in Lk neurons abolished the suppression of feeding induced by optogenetic activation. **B** (right), RNAi in Lk neurons of analogous receptors for other candidate neuromodulators had no effect. Adjacent schematic shows that Lk neurons convey hunger state information to downstream neurons expressing Lk receptors. Lk neuron activation suppresses feeding, conveying decreased hunger, while Lk inhibition leads to consumption of higher caloric food, conveying increased hunger. **C** (left), Optogenetic activation of Allatostatin A (AstA) neurons shifted preference towards sweet while optogenetic inhibition shifted it towards bittersweet. Simultaneous AstA RNAi abolished the preference shift induced by optogenetic activation. **C** (right), Dop1R1 RNAi in AstA neurons shifted preference towards sweet. Adjacent schematic shows that AstA neurons receive decision-relevant direct dopaminergic inputs through Dop1R1. Both AstA neuron activation and inhibition shift food decision in opposite directions and send decision-relevant information to downstream neurons expressing AstA receptors. **D** (left), Optogenetic activation of NPF neurons shifted preference towards sweet. Simultaneous NPF RNAi abolished the preference shift induced by optogenetic activation. **D** (right), Lkr or Dop1R1 RNAi in NPF neurons each shifted preference towards sweet. Adjacent schematic shows that NPF neurons receive decision-relevant direct dopaminergic inputs through Dop1R1 and Lk inputs through Lkr receptors. NPF neuron activation shifts food decision to sweet and NPF neurons send decision-relevant information to downstream neurons expressing NPF receptors. **E** (left), Optogenetic activation of DH44 neurons had no effect on preference, while optogenetic inhibition shifted preference towards bittersweet. **E** (right), DopEcR RNAi in DH44 neurons shifted preference towards bittersweet. Adjacent schematic shows that DH44 neurons receive decision-relevant direct dopaminergic inputs through DopEcR. DH44 neuron inhibition shifts food decision to bittersweet and DH44 neurons send decision-relevant information to downstream neurons expressing DH44 receptors. Plots show mean ± 95% CI, with violins depicting full data distribution; 5 ≤ trials ≤ 30 per violin, mode = 10. See Table S1 for sample size and statistics. p<0.00001=****, p<0.0001=***, p<0.01=**, p<0.05=*.

The specific effects of activation or inhibition of restricted neuron populations on the sweet versus bittersweet decision provides insights into the roles of such populations in value estimation and integration. Optogenetic activation of Leucokinin (Lk) neurons suppressed feeding in food deprived flies (Fig. 2A, 2B left panel, Fig. S2A-B), suggesting that Lk may encode or relay metabolic state information. To confirm that Lk secreted by these neurons was the molecular basis of this feeding suppression, we simultaneously knocked down Lk expression with genetically encoded RNAi while optogenetically activating Lk neurons. While almost no flies consumed food when Lk neurons were activated without Lk RNAi, simultaneous Lk knockdown prevented this feeding suppression (Fig. 2B left panel, Fig. S2A-B). This indicates that Lk secretion mediates feeding suppression by Lk neurons. Optogenetic silencing of Lk neurons shifted preference towards bittersweet (Fig. 2A, 2B left panel, Fig. S2A-B). Feeding suppression induced by Lk neuron activation implies a decrease in perceived hunger of food deprived flies, consistent with the shift towards higher calorie bittersweet food by Lk neuron inhibition reflecting increased perceived hunger. Optogenetic activation of Allatostatin A (AstA) neurons shifted preference towards sweet, while inhibition shifted preference towards bittersweet (Fig. 2A, 2C left panel). We confirmed that AstA was the molecular basis of this shift in preference by simultaneous AstA RNAi knockdown and optogenetic activation of AstA neurons (Fig. 2C left panel). Optogenetic activation of NPF neurons shifted preference towards sweet, and this shift was abolished by simultaneous activation and NPF RNAi knockdown (Fig. 2A, 2D left panel). Optogenetic activation of DH44 neurons had no significant effect, but inhibition shifted preference towards bittersweet (Fig. 2A, 2E left). Dopaminergic subsets involved in aversive memory (Fig. 2A PPL1-γ2α’1) (Berry et al., 2012), taste conditioning (Fig. 2A PPL1- α3) (Masek et al., 2015), and long-term memory (Fig. 2A PAM-α1) (Yamagata et al., 2015) also affected food choice. Specifically, optogenetic activation of these dopaminergic subsets shifted preference towards bittersweet (Fig. 2A). Optogenetic activation of various subsets of mushroom body neurons, a brain region controlling higher-order behaviors, did not affect preference (Fig. 2A). However, optogenetic inhibition of a specific subset of FBl6 neurons shifted preference towards bittersweet (Fig. 2A, 3A left panel). Value estimates of internal state and external sensory cues, which are likely computed by identified modulatory neurons, are crucial for decision making. Because many of these neuromodulators and their receptors (Al-Anzi et al., 2010; Cavey et al., 2016; Donlea et al., 2018; Kahsai and Winther, 2011; Kim et al., 2020; Qian et al., 2017) co-localize in the fan-shaped body, we thus hypothesize that neuromodulatory inputs encoding value estimates are integrated in FBl6, where a decision is generated.

**Figure 3.**
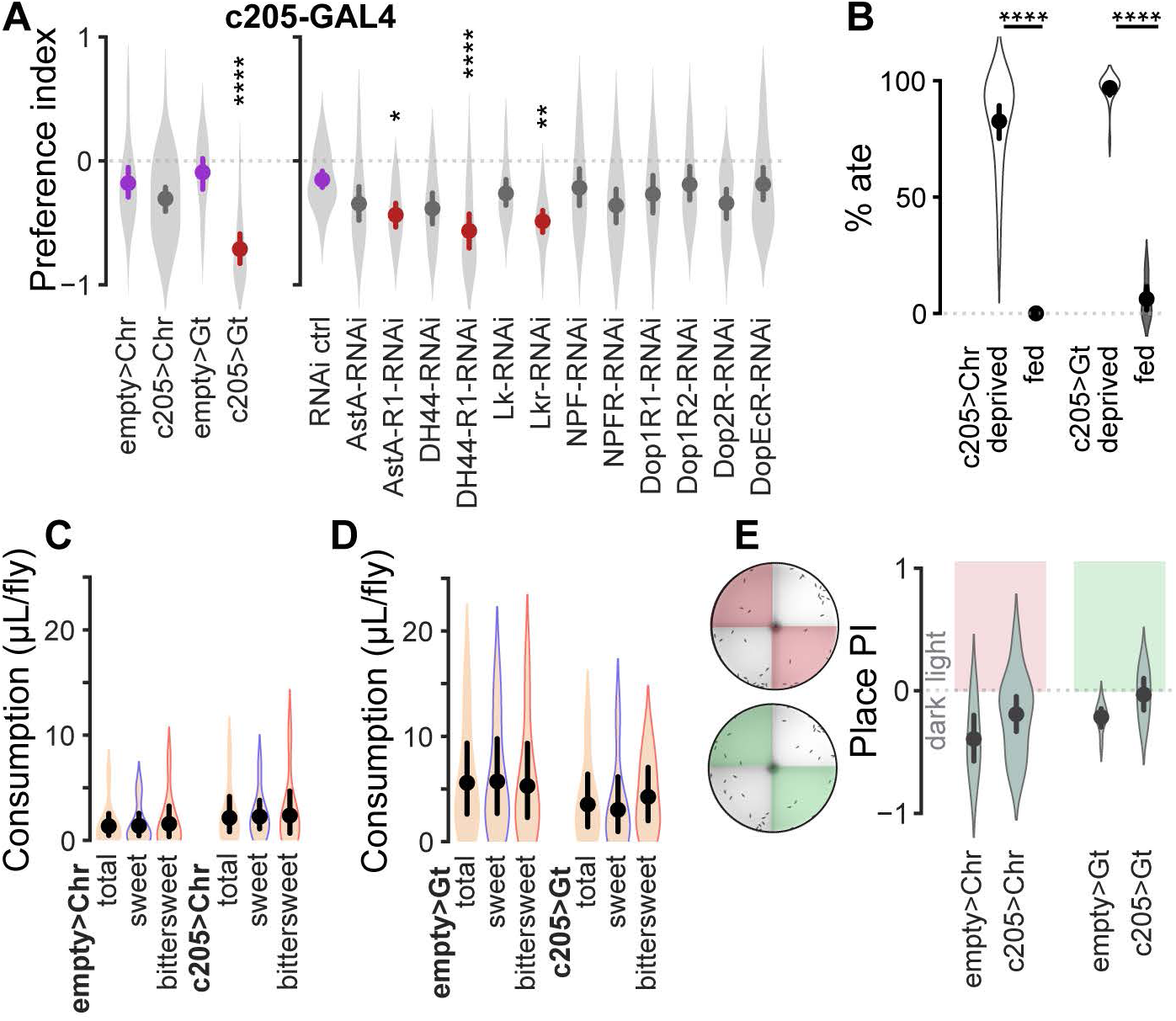
Fan-shaped Body layer 6 is the convergence node of a decision making ensemble. **A** (left), Optogenetic activation of FBl6 neurons did not affect preference. In contrast, optogenetic inhibition shifted preference towards bittersweet. **A** (right), RNAi knockdown of AstA-R1, DH44-R1, or Lkr receptors in FBl6 shifted preference towards bittersweet. **B**, Optogenetic activation or inhibition of FBl6 did not induce feeding in fed flies or prevent feeding in food-deprived flies. **C-D**, Optogenetic activation (**C**) or inhibition (**D**) of FBl6 did not change total food consumption per fly compared to respective empty-GAL4 controls. Food consumption was quantified by extracting red and blue food dyes from flies after the assay and measuring the extract absorbance using UV-vis spectrometry. Concentrations of individual dyes were calculated per trial by interpolating absorbance in the standard curve for each dye. Total volume of agarose ingested was then calculated from red and blue dye concentrations and divided by number of flies that ate red and blue food in that trial. Total food volume is the average of consumption by each fly regardless of type of food consumed. Within each group, there was no significant difference in sweet vs. bittersweet food consumption per fly. Optogenetic inhibition of FBl6 shifted preference towards bittersweet (**A,** left), because more flies consumed bittersweet than sweet. **E**, Optogenetic activation or inhibition of FBl6 did not significantly change preference (place PI) of flies for illuminated vs. non-illuminated sectors of the arena, indicating that neither activation nor inhibition of FBl6 is inherently rewarding or punishing. Plots show mean ± 95% CI, with violins depicting full data distribution. Statistically different means are shown in different color as in Fig. 2. See Table S1 for sample size and statistics. p<0.00001=****, p<0.0001=***, p<0.01=**, p<0.05=*.

To test this hypothesis and determine whether the neurons we identified in our optogenetic screen are connected in a behaviorally relevant ensemble, we employed a chemoconnectomics approach (Deng et al., 2019) using cell-specific genetically encoded RNAi knockdown of neuropeptide and dopamine receptors. By knocking down a specific receptor in a candidate target neuron, we test whether that neuron receives direct input from a neuron secreting the cognate ligand for that receptor. Knockdown of neuropeptide or dopamine receptors in Lk neurons did not shift preference (Fig. 2B right panel). This indicates that Lk neurons do not receive direct inputs from decision-relevant dopaminergic neurons or from neurons secreting the ligands for tested neuropeptide receptors, and implies that Lk neurons must receive food preference and hunger related information indirectly from other neurons. RNAi knockdown of Dop1R1 dopamine receptor in AstA neurons shifted preference towards sweet (Fig. 2C right panel), indicating that AstA neurons receive food-preference-relevant dopaminergic inputs. RNAi knockdown of Lkr and Dop1R1 in NPF neurons shifted preference towards sweet (Fig. 2D right panel), indicating that NPF neurons receive food-preference-relevant Lk and dopaminergic inputs. RNAi knockdown of DopEcR dopamine receptor in DH44 neurons shifted preference towards bittersweet (Fig. 2E right panel), indicating that DH44 neurons receive food-preference-relevant dopaminergic inputs.

Importantly, RNAi knockdown of Lkr, AstA-R1, or DH44-R1 receptors in FBl6 neurons shifted preference towards bittersweet (Fig. 3A right panel), indicating that FBl6 neurons are directly modulated by these three neuropeptides to affect food choice. Furthermore, direction of change in food preference induced by receptor RNAi in FBl6 mirrors direction of change in food preference induced by optogenetic inhibition of the corresponding neuropeptide neurons. First, both AstA-R1 receptor RNAi in FBl6 neurons (Fig. 3A right panel) and optogenetic inhibition of AstA neurons (Fig. 2C left panel) shifted food preference towards bittersweet. Second, both DH44-R1 receptor RNAi in FBl6 (Fig. 3A right panel) and optogenetic inhibition of DH44 neurons (Fig. 2D right panel) shifted preference towards bittersweet. Finally, both Lkr receptor RNAi in FBl6 (Fig. 3A right panel) and optogenetic inhibition of Lk neurons (Fig. 2B right panel) shifted preference towards bittersweet. RNAi knockdown of dopamine receptors in FBl6 had no effect (Fig. 3A right panel). This matrixed chemoconnectomics strategy mapped the neuromodulatory connections between nodes in the ensemble to control choice, and uncovered FBl6 as a previously unknown convergence node that is well positioned to integrate sensory, metabolic, and experiential information for decision making.

### Fan-shaped body neurons encode choice

Value estimates of internal state variables, such as hunger, external environmental cues, appetitive or aversive value of food (i.e., valence), and past sensorimotor experience are integrated with one another and transformed into behavioral choice. This raises the key question whether FBl6 neurons compute value estimates or, instead, integrate these value estimates to encode choice. If FBl6 neurons encode or estimate value of metabolic parameters such as hunger or satiety, manipulating their activity would be expected to directly influence feeding behavior. To test this, we first quantified the proportion of flies that consumed food during optogenetic activation or inhibition of FBl6 neurons. During FBl6 neural activity manipulation, most food-deprived flies consumed food while most fed flies did not (Fig. 3B, Table S1), revealing that FBl6 activity manipulation neither prevented food consumption by food-deprived flies nor induced food consumption by fed flies. These results demonstrate that hunger state is not affected by FBl6 neural activity. Next, we quantified food consumption per fly during optogenetic activation or inhibition of FBl6 neurons and found no effect on total amount of food consumed per fly (Fig. 3C, Table S1). These manipulations also did not affect the quantity of sweet versus bittersweet food consumed per fly within each condition (Fig. 3D, Table S1), again demonstrating that FBl6 neural activity does not affect hunger state. Rather, the shift in food preference induced by optogenetic inhibition of FBl6 (Fig. 3A left panel) was due to an increase in the proportion of flies consuming bittersweet over sweet food, and not each fly consuming a larger quantity of bittersweet food. Taken together, these results demonstrate that FBl6 does not encode or affect metabolic signals of hunger or satiety.

If a neural population encodes reward or punishment, then manipulating the activity of that population could directly shift a decision without integrating value estimates or encoding choice. To test whether FBl6 activity is intrinsically rewarding or punishing, we illuminated half the decision arena with either red (for optogenetic activation) or green (for optogenetic inhibition) light in the absence of food and quantified preference for the illuminated versus dark sectors. Flies expressing optogenetic actuators CsChrimson or GtACR1 in FBl6 did not alter their preference for illuminated versus dark sectors compared to non-expressing controls (Fig. 3E), demonstrating that FBl6 activation or inhibition is intrinsically neither rewarding nor punishing. Animals accumulate information about past experience to inform future decisions. We hypothesized that FBl6 integrates internal hunger state and external food-related value estimates with experiential information to drive decisions. To understand how past experience affects FBl6 activity, we recorded FBl6 neural taste responses in flies that had different food-related experiences. Equal-preference taste stimuli were presented to forelegs of the fly (Fig. 4A) while measuring intracellular Ca^2+^ signals ratiometrically using GCaMP6f (Chen et al., 2013) and tdTomato (Fig. 4B) expressed in FBl6 (Fig. 4B-C). First, we tested the effect of hunger on FBl6 taste responses in naïve flies, i.e., flies that had not experienced the decision task at all. FBl6 neurons of naïve food-deprived flies were strongly inhibited by the bittersweet stimulus, but not sweet (Fig. 4D-E, “naïve deprived”). Flies often find bittersweet food aversive (Masek and Scott, 2010), and thus inhibitory taste responses in FBl6 to bittersweet food may represent rejection of this option. In contrast to hungry food-deprived flies, fed flies reject both foods in the decision task (Fig. 3B). If FBl6 neural activity represents rejected choice, then we expect FBl6 inhibition by both sweet and bittersweet stimuli in naïve fed flies. Indeed, FBl6 neurons of naïve fed flies were strongly inhibited by both bittersweet and sweet stimuli (Fig. 4D-E, “naïve fed”). Importantly, this is the first observation of gustatory responses in the *Drosophila* fan-shaped body, or any other part of the central complex.

**Figure 4.**
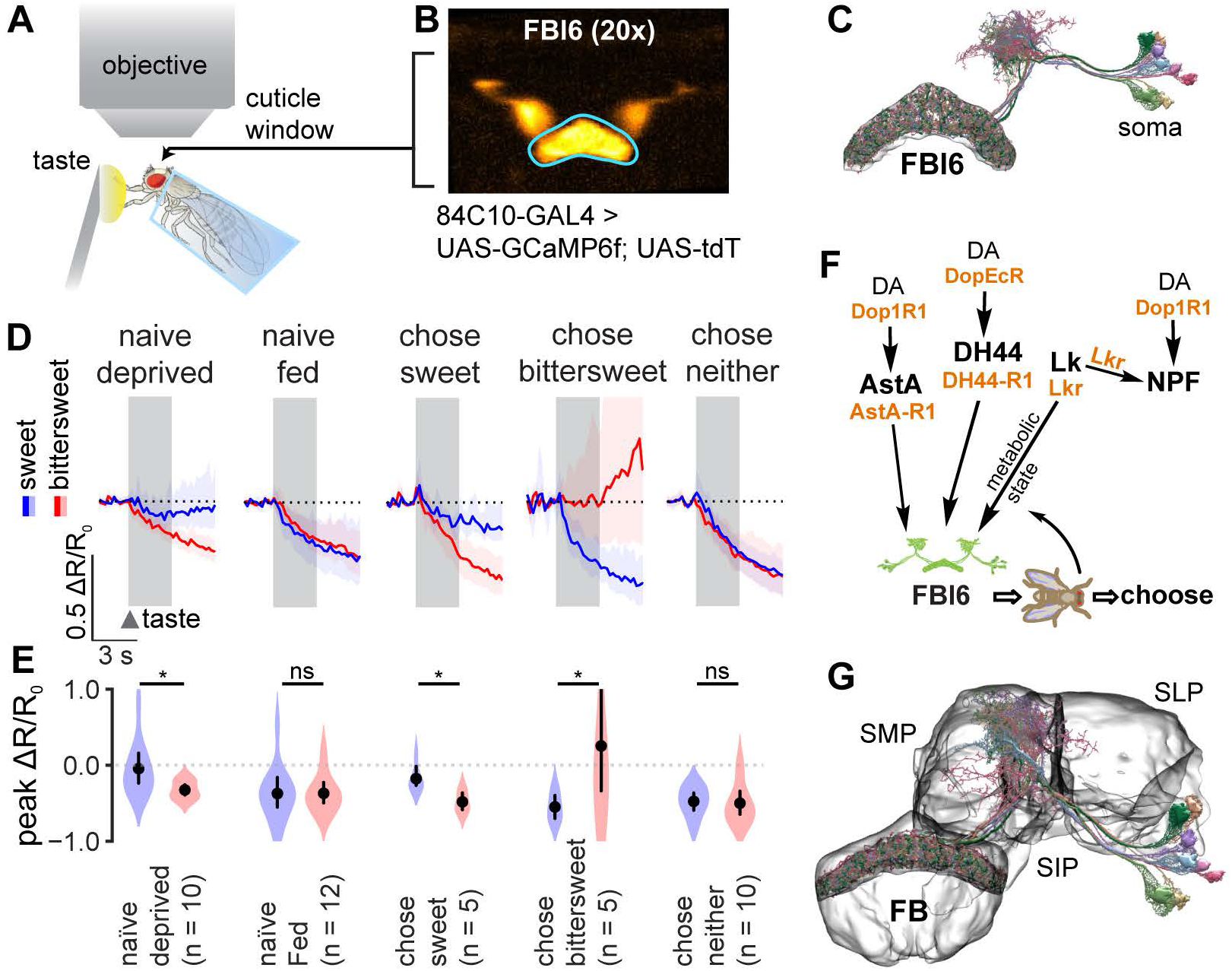
Neural activity in FBl6 encodes food choice. **A**, *In vivo* Ca^2+^ imaging of taste responses in FBl6 neurons of flies in different hunger states making different sweet-bittersweet decisions. Sweet or bittersweet taste stimuli are applied to the forelegs, and changes in Ca^2+^ responses are measured ratiometrically in FBl6 using GCaMP6f and tdTomato. **B,** Neuronal expression in FBl6 reported by tdTomato. Region of interest for fluorescence measurement is outlined in cyan. 84C10-GAL4 driver line is used to specifically target strong fluorescent protein expression in FBl6 neurons (FlyLight; Jenett et al., 2012) (Fig. S3B-D). **C,** EM reconstruction of FBl6 neurons shows neural projections restricted to layer 6 of the fan-shaped body. EM reconstruction of traced FBl6 neurons from the Janelia hemibrain data was performed using neuPrint and NeuronBridge for neurons targeted by 84C10-GAL4 and embedded in the surface representation of standardized FBl6 brain region. **D,** Ratiometric calcium responses, ΔR/R_0_, of flies with different hunger state and decision outcomes. Sweet (50mM sucrose) and bittersweet (500 mM sucrose + 1 mM quinine) taste stimuli from the equal-preference condition were applied for 3 s, and neural response was quantified for 4 s post-stimulus application. Taste stimulus application is indicated by gray background region. FBl6 neurons were strongly inhibited by the behaviorally rejected taste stimulus. Specifically, FBl6 neurons of flies that chose sweet were inhibited by bittersweet taste, while FBl6 neurons of flies that chose bittersweet were inhibited by sweet taste. (**D-E**). Calcium activity trace depicts mean ΔR/R_0_ ± 95% CI. **E**, Peak ΔR/R_0_ shows significant difference between responses to rejected vs. chosen taste within each fly condition. p<0.05=* (see Table S1 for details on statistics). Points on graphs depict mean ± 95% CI, with violins depicting full data distribution. **F,** Schematic of the decision making ensemble converging on FBl6. FBl6 activity is the neural representation of behavioral food choice. This activity is modulated by taste, previous experience, and hunger state. FBl6 neurons likely receive inputs encoding these parameters directly through AstA, DH44, and Lk receptor signaling, and indirectly through NPF and dopamine (DA) pathways. FBl6 integrates the converging information to form a representation of choice, which is relayed to downstream motor circuits for behavior implementation. **G**, EM reconstruction of FBl6 neurons targeted by 84C10-GAL4 was performed as previously and embedded in the surface representation of standardized whole fan-shaped body (FB). FBl6 neural projections to superior medial protocerebrum (SMP), superior intermediate protocerebrum (SIP), and superior lateral protocerebrum (SLP) higher brain regions are shown embedded in the surface representation of these standardized higher brain regions.

If, as in naïve flies, FBl6 neural activity also represents rejected choice in flies that have had the opportunity to decide between sweet and bittersweet, then we expect decision outcomes to affect FBl6 taste responses. Specifically, we expect FBl6 neurons of each fly to be strongly inhibited by the rejected choice for that fly, based on its individual decision. Consistent with this, when taste responses were subsequently measured in flies that chose sweet food in the decision task, FBl6 neurons were strongly inhibited by the rejected bittersweet stimulus, but not by the chosen sweet (Fig. 4D-E, “chose sweet”). Furthermore, FBl6 neurons of flies that chose bittersweet food were strongly inhibited by the rejected sweet stimulus, but not by the chosen bittersweet (Fig. 4D-E, “chose bittersweet”). Finally, FBl6 neurons of flies that chose neither option, i.e., rejected both, were strongly inhibited by both bittersweet and sweet stimuli (Fig. 4D-E, “chose neither”). In summary, FBl6 neural activity is strongly inhibited by the taste of food that each individual fly rejects, suggesting that suppression of FBl6 activity is the neural representation of behavioral food choice.

We then tested whether FBl6 neurons encode not only taste of rejected food, but also behavioral choice associated with other sensory cues. Specifically, we determined whether FBl6 responds to water, and whether such responses are dependent on internal state and prior experience of each fly. While FB16 neurons of food-deprived flies that chose sweet or neither in the decision assay were inhibited by water applied to the forelegs, those of fed or food-deprived naive flies were not (Fig. S4A-B). Furthermore, choice outcome in the decision assay affected water responses, with water inhibiting FB16 neurons of flies choosing sweet or neither, but not of flies choosing bittersweet (Fig. S4A-B). Individual flies that chose bittersweet higher-calorie food over sweet are likely hungrier based on their individual behaviors during maintenance and food deprivation than flies that chose lower-calorie sweet or neither (i.e., didn’t consume), and may correspondingly be thirstier. This could then suggest that the FBl6 inhibition by water we observed specifically in flies that chose sweet or neither represents rejection of water by these less-thirsty flies, and that lack of inhibition in flies that chose bittersweet represents acceptance. Regardless of the specific interpretation (which requires further study), these water responses indicate that, like for taste, FBl6 encodes not stimulus identity *per se*, but rather state- and experience-dependent behavioral choice.

These state- and experience-dependent FBl6 responses do not simply encode taste or water identity, but rather encode behavioral choice that arises from integration of internal state and external sensory inputs. This neural representation of choice is modulated by taste (sweet vs. bittersweet), previous experience (naïve vs. experience with the two-choice task, including decision outcome), and hunger state (food-deprived vs. fed) (Fig. 4D-E, Fig. S4A-B). FBl6 neurons likely receive inputs encoding these parameters directly from AstA, DH44, and Lk neurons, and indirectly from NPF and dopamine neurons. These signals within the decision ensemble are then integrated by FBl6 to generate a representation of choice, which is relayed to downstream motor circuits to implement the behavioral decision (Fig. 4F).

## Discussion

Animals make decisions about which foods to consume by integrating internal physiological state signals with external sensory cues. Here we delineated a neuronal ensemble in *Drosophila* that underlies food-related decision making during sensory conflict between sweet and bittersweet food choices (Fig. 1). Activating or silencing particular nodes in this ensemble shifts the decision balance between sweet and bittersweet choices (Fig. 2, 3). Inputs encoding hunger state, taste identity, and past experience converge on FBl6 neurons, which integrate these inputs to generate a neural representation of food choice that drives motor outputs for foraging decisions (Fig. 4).

Animals assess and assign value estimates to internal state and external environmental parameters before integrating these estimates to guide adaptive decision making. Upstream neuromodulatory subsets in the decision ensemble we identified have known roles in hunger dependent food intake (Dus et al., 2015; Hentze et al., 2015; Yang et al., 2018; Zandawala et al., 2018), reward (Shao et al., 2017), valence (Berry et al., 2012; Masek et al., 2015), and long-term memory (Berry et al., 2012; Masek et al., 2015; Yamagata et al., 2015). These modulatory neurons are well positioned to estimate value of features in the sensory environment and internal hunger state. For example, AstA neuron activity influences relative carbohydrate and protein preference (Hentze et al., 2015), while DH44 neurons sense sugars (Dus et al., 2015) and amino acids (Yang et al., 2018). Thus, AstA and DH44 neurons could convey food identity information to FBl6. NPF neuron activation is inherently rewarding (Shao et al., 2017), and thus could convey food valence information. Lk neurons have been implicated in nutrient sensing (Zandawala et al., 2018), and their activity regulates feeding in food deprived flies (Fig. 2A-B), suggesting that internal metabolic state information could reach FBl6 through Lk/Lkr signaling. Dopaminergic subsets involved in aversive memory (PPL1 γ2α’1) (Berry et al., 2012), taste conditioning (PPL1 α3) (Masek et al., 2015), and long-term memory (PAM α1) (Yamagata et al., 2015) also affected food choice (Fig. 2A) and could provide signals for predicting and updating value estimates in working memory, analogously to primate dopaminergic ventral tegmental area (Sugrue et al., 2005). FBl6 neurons project their axons into the fan-shaped body (Kim et al., 2020; Qian et al., 2017), have dense dendritic projections in the superior medial protocerebrum (SMP), and have sparse dendritic projections in superior intermediate protocerebrum (SIP) and superior lateral protocerebrum (SLP) (Kim et al., 2020; Qian et al., 2017) (Fig. 4G). In these higher brain regions, FBl6 receives synaptic input from dopaminergic neurons (Liu et al., 2012b; Pimentel et al., 2016) that regulate sleep (Liu et al., 2012b; Pimentel et al., 2016) and ethanol preference (Azanchi et al., 2013). Interestingly, direct dopaminergic input to FBl6 through dopamine receptors did not influence food choice (Fig. 3A). Instead, indirect dopaminergic inputs likely conveyed by AstA, DH44, and NPF neurons regulated food choice (Fig. 2B-E).

Other neuromodulatory neurons, second-order sensory projection neurons, and interneurons also likely interact with this decision ensemble to convey gustatory and other sensory inputs. There are no known projections of primary gustatory sensory neurons or second-order taste neurons (Bohra et al., 2018; Kain and Dahanukar, 2015; Kim et al., 2017a; Miyazaki et al., 2015) to the fan-shaped body. Food preference was affected neither by acute optogenetic nor by chronic inhibition (using tetanus toxin light chain to impair vesicle docking) of the only known taste projection neurons (TPN3) that relay bitter taste information to SLP and are essential for conditioned taste aversion (Kim et al., 2017a) (Fig. S5A). However, SMP, SLP, and SIP higher brain regions, to which FBl6 neurons also densely project (Fig. 4G), have been implicated as target areas for second-order taste neurons (Talay et al., 2017). It thus is possible that sweet and bitter taste information reaches the fan-shaped body through second-order taste projection neurons that terminate in these higher brain regions. Another possibility is that taste identity information is conveyed to FBl6 by the dopaminergic neurons that respond to sweet (Huetteroth et al., 2015) and bitter (Masek et al., 2015) tastes, are involved in taste conditioning (Huetteroth et al., 2015; Masek et al., 2015), and shift preference in our decision assay (Fig. 2A, PAM-α1, PPL1-α3).

A value integrator for food-related decision making requires as inputs estimates of taste identity, previous experience, and hunger state. FBl6 neuron activity is modulated by these parameters and strongly inhibited by the taste of rejected food choice. Lack of FBl6 inhibition in response to a taste stimulus could represent acceptance of that option (Fig. 4D-E). The mechanistic connection between optogenetic inhibition of FBl6 shifting food preference to bittersweet (Fig. 3A), on the one hand, and rejected food choice strongly inhibiting FBl6 neural activity (Fig. 4D-E), on the other hand, requires further investigation. It is possible that optogenetic inhibition of FBl6 throughout the decision task disrupts value integration and decision making resulting in the hungry fly defaulting to higher calorie bittersweet food. Regardless, our findings indicate that the behavioral response of an individual fly can be predicted by dynamic neural activity in FBl6.

Mammalian studies provide converging evidence for multiple interconnected networks in frontal cortex and basal ganglia that compute and store value estimates of sensory environment and motor events in that environment required for decision making (Lee et al., 2012; Sugrue et al., 2005). The neural ensemble described in this study is an analogous framework of interconnected networks for potentially storing, computing, and updating value estimates that are likely integrated by FBl6 neurons. While decision making theories in mammals have focused on how values are represented in the brain (Lee et al., 2012; Sugrue et al., 2005), mechanisms by which the brain integrates value information to make decisions remain unclear (Shadlen and Kiani, 2013; Tsetsos et al., 2012). Changes in neural spiking pattern are thought to underlie sensory information accumulation and integration, with a decision being made when neural firing rate reaches a threshold (Gold and Shadlen, 2007; Park et al., 2014; Wang, 2008). Persistent neural activity and synaptic plasticity changes are thought to underlie storage of sensory information and past choice history to guide adaptive decisions (Wang, 2008). Future work is required to test the proposed hypotheses of specific roles for each node in the fly decision ensemble, how upstream inputs are integrated in FBl6, how this integration is transformed into the representation of choice, and how downstream motor circuits implement the foraging decision.

## Methods

### Fly husbandry

Flies were cultured on standard cornmeal medium on 12:12 light:dark cycle at 25⁰C. *w1118* lab stock was used as wild type. All other genotypes and their sources are described in Table S2. 2-5 day old flies were wet starved for 2-48 h (based on experiment design) on a wet Kimwipe with 1.5 ml distilled water. For optogenetic experiments, flies were food deprived for 21 h before testing on 0.4 mM all-*trans* Retinal (Cayman Chemicals) in 1% agar. Flies for RNAi knockdown and their controls were moved to 28⁰C for 21 h the day before testing, i.e., during food deprivation, to induce strong RNAi. RNAi control was created for each GAL4 line by crossing the respective GAL4 to UAS-Valium (see Table S2). All RNAi lines that we used were from Harvard TRiP project (Ni et al., 2009; Perkins et al., 2015) and have been validated by independent groups (see Table S2). Flies for simultaneous optogenetic and RNAi experiments were created using the genotypes mentioned in Table S2. All experiments were conducted at Zeitgeber Time 3-6.

### Two-choice assay and optogenetics

Sweet foods were made with different concentrations of sucrose and bittersweet foods with 500mM sucrose (Sigma) and 1mM quinine (Alfa Aesar or Beantown Chemicals, CAS#207671-44-1) dissolved in 1% agarose (AmericanBio) made in distilled water. 0.04% w/v red dye (Sulforhodamine B, MP Biochemicals, CAS# 3520-42-1) and 0.02% w/v blue dye (Erioglaucine A, Alfa Aesar, CAS# 3844-45-9) were used for food coloring. Dye colors were alternated between sweet and bittersweet foods for each condition and there was no preference for one dye over the other at the concentrations used. Fly arenas were prepared by pouring agarose based foods in two-compartment petri-dishes (90-100 mm diameter) from Kord Valmark, EMS, or Fisher Scientific. Because of a thin physical barrier between the two compartments in the arena there was no diffusion between the two foods. Groups of 20-35 flies were aspirated and introduced into the arena 5-10 sec before the start of the experiment. All experiments were conducted in dark so that there was no effect of food color on preference. Arenas were placed on a platform with IR backlight for video recording using a Flea Pointgrey camera (FL3-U3-20E4C/M) at 15 fps. For optogenetics, we used high-power LEDs (Luxeon) placed adjacent to backlight IR LEDs (based on Janelia ID&F design) of 627 nm (for CsChrimson) and 520 nm (for GtACR1) that were controlled using Arduino Uno. For optogenetic screen, both red and green lights were pulsed at 100% max intensity, 50Hz, 25% duty cycle. For follow up experiments, CsChrimson experiments were conducted at 25% max intensity; GtACR1 follow up was done at screen condition. Light was pulsed for the entire duration of the experiment. At the end of the experiment, flies were anaesthetized using CO2 and their belly color was recorded under a dissection microscope. Preference index (PI) was calculated as (no. of sweet food flies + 0.5 no. of both food flies) - (no. of bittersweet food flies + 0.5 no. of both food flies) / no. of total flies that ate, where negative PI would mean that more number of flies ate bittersweet food.

### Food intake quantification

Food intake was quantified using spectrophotometry as previously described (Deshpande et al., 2014; Wong et al., 2009). After recording belly color flies were frozen in 1.5 ml Eppendorf tubes at -20⁰C until intake quantification (1-2 days). Flies from each trial were separately homogenized in distilled water (5 µl /fly) using a motorized pestle (BT Labsystems, BT703) for 1.5 min and centrifuged at 13000 rpm for 5 min. Absorbance of the debris-cleared 2 µl supernatant was measured on NanoDrop 2000 Spectrophotometer (Thermo Fisher Scientific) at 565 nm (for red dye) and 630nm (for blue dye). Flies that ate uncolored 1% agarose with 50mM sucrose were used as blank for baseline control. Red and blue dye concentrations were interpolated using their respective standard curves (GraphPad Prism) acquired from serial dilutions of single dyes in distilled water. Since we knew the number of flies that ate each color per trial, we could calculate per fly blue and red concentrations in the same solution.

### Calcium imaging and data analysis

3-5 day old flies (naïve or after two-choice assay) were aspirated and positioned in a custom made fly holder in which they were glued using two-part transparent epoxy (Devcon). Only the top of fly head (for imaging) and the forelegs (for taste delivery) were outside the holder, while the rest of the fly, including proboscis were restrained in the fly holder. No anesthesia was used. A small piece of head cuticle was dissected and air sacs removed using a 30-gauge syringe needle and fine forceps, immediately followed by sealing the head capsule with a translucent surgical silicone adhesive (Kwil-Sil, WPI). Dissected fly was then placed in a humidified chamber for 15 min recovery before imaging.

Calcium imaging was performed on a Zeiss Axio Examiner upright microscope with 20x air objective and a Colibri module for LED control. tdTomato was excited at 555 nm (80% intensity) and GCaMP6 at 470 nm (100% intensity). An image splitter (Photometrics DV-2) was used to split red and green channels and acquire simultaneous images for tdTomato and GCaMP6 using a Hamamatsu ORCA-R2 C10600 camera. Images were acquired at 5 fps with variable baseline (3 to 10 s required for stable tastant delivery) followed by tastant application to the forelegs, using a syringe, for 3 s and 4 s of no tastant. Excess tastant was wicked from the forelegs using absorbent tissue paper between each application. 10 s inter-trial interval was used during which all lights were off. Water was always applied first, followed by either sweet or bittersweet tastant. Sequence of sweet and bittersweet was alternated between flies. Sweet: 50 mM sucrose in distilled water; bittersweet: 500 mM sucrose + 1 mM quinine in distilled water. For flies that chose sweet in the two-choice behavior assay only trials with sweet as the first tastant were averaged and for flies that chose bittersweet, only trials with bittersweet as the first tastant were averaged.

Pixel intensities were extracted in Fiji followed by data analysis in MATLAB, both using custom written code. After background subtraction using Fiji’s rolling-ball method (20 px), ROIs were manually drawn and saved on the tdTomato image, and superimposed on the GCaMP image (both reporters were expressed in the same neurons using the same driver) for mean ROI pixel intensity extraction. The saved intensity signals were then analyzed in MATLAB. tdTomato and GCaMP traces were individually corrected for photobleaching by fitting a single exponential function. Corrected GCaMP trace was then divided by the corrected tdTomato trace to obtain the ratiometric fluorescence trace (R). For relative fluorescence fold change (ΔR/R_0_) determination, baseline fluorescence (R0) was calculated by averaging R over 2 s preceding tastant application. Peak ΔR/R_0_ was calculated during 4 s following tastant application.

### EM reconstruction

Electron microscopy images were reconstructed from publically available Janelia FlyEM hemibrain data using neuPRINT (Xu et al., 2020). Neuron identities were confirmed in NeuronBridge (Jody et al., 2020) by cross-referencing EM traced FBl6 neurons matched with light microscopy images of 84C10-GAL4 from FlyLight (Meissner et al., 2020). FBl6 mesh, whole FB mesh, and SMP, SIP and SLP brain region meshes were used to depict brain regions with neural projection areas.

### Statistics

All data were plotted in either Python or MATLAB using custom written code. Statistics were carried out in GraphPad Prism. If all data passed Kolmogorov-Smirnov normality test, ANOVA was conducted, otherwise Kruskal-Wallis test was conducted, both followed by appropriate post-hoc tests. Details of statistics for each figure are provided in Table S1. Sample sizes are reported in parentheses next to dataset name in Table S1.

## Acknowledgements

We thank Peter Niesman for help with calcium imaging data collection and Jason Braco for helpful discussions. Yichen Luo from John Carlson’s lab provided useful information for design of the taste delivery imaging rig. Tanya Wolff provided insights into fan-shaped body neuroanatomy. We also thank Gerry Rubin and Tanya Wolff for sharing unpublished fly lines ss00208 and ss00225. These studies were supported in part by the National Institute of Neurological Disorder and Stroke, National Institutes of Health (NIH) (R01NS091070) and the National Institute of General Medical Sciences, NIH (R01GM098932).

## Author contributions

P.S. conceptualized the study, designed and performed experiments, and analyzed data; L.Y.M. contributed to experimental design and fly dissection; P.S. and M.N.N. interpreted data; P.S. and M.N.N. prepared the manuscript with input from all authors.

## Data and code availability

Further information and requests for resources and reagents should be directed to and will be fulfilled by the Lead Contact, Dr. Michael N. Nitabach. The data and custom code that support the findings from this study are available from the Lead Contact upon request.

---

**Table S1.**
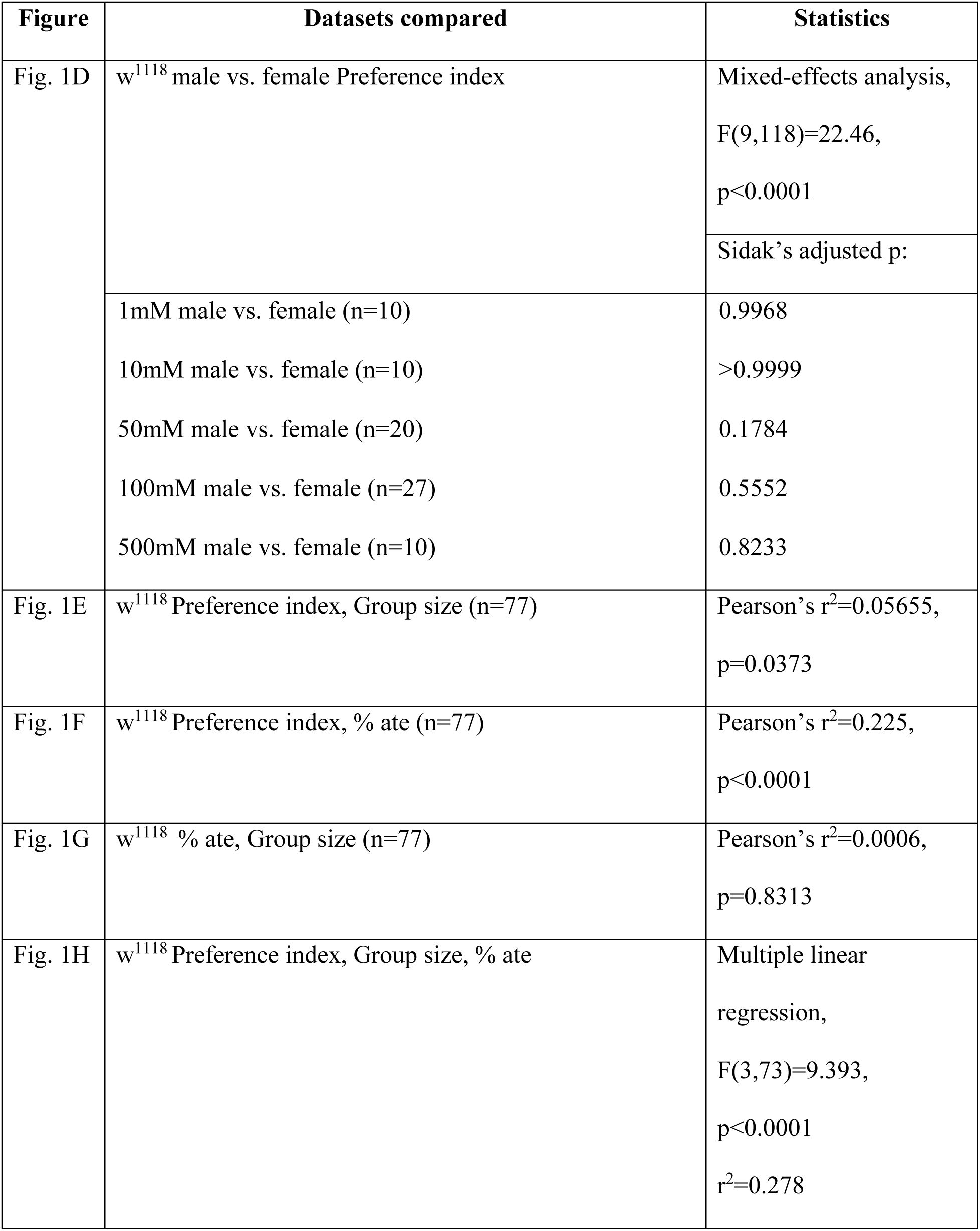
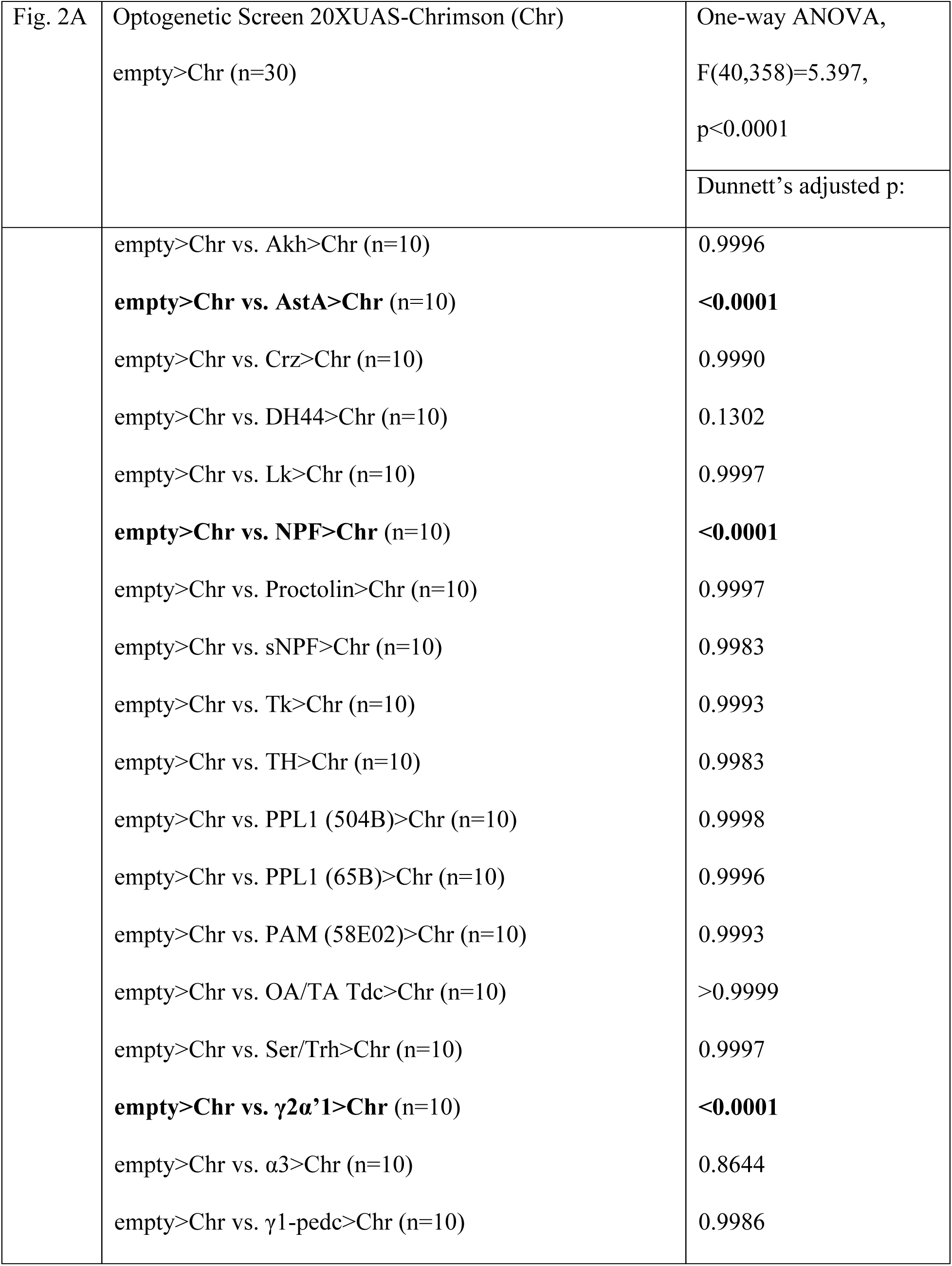
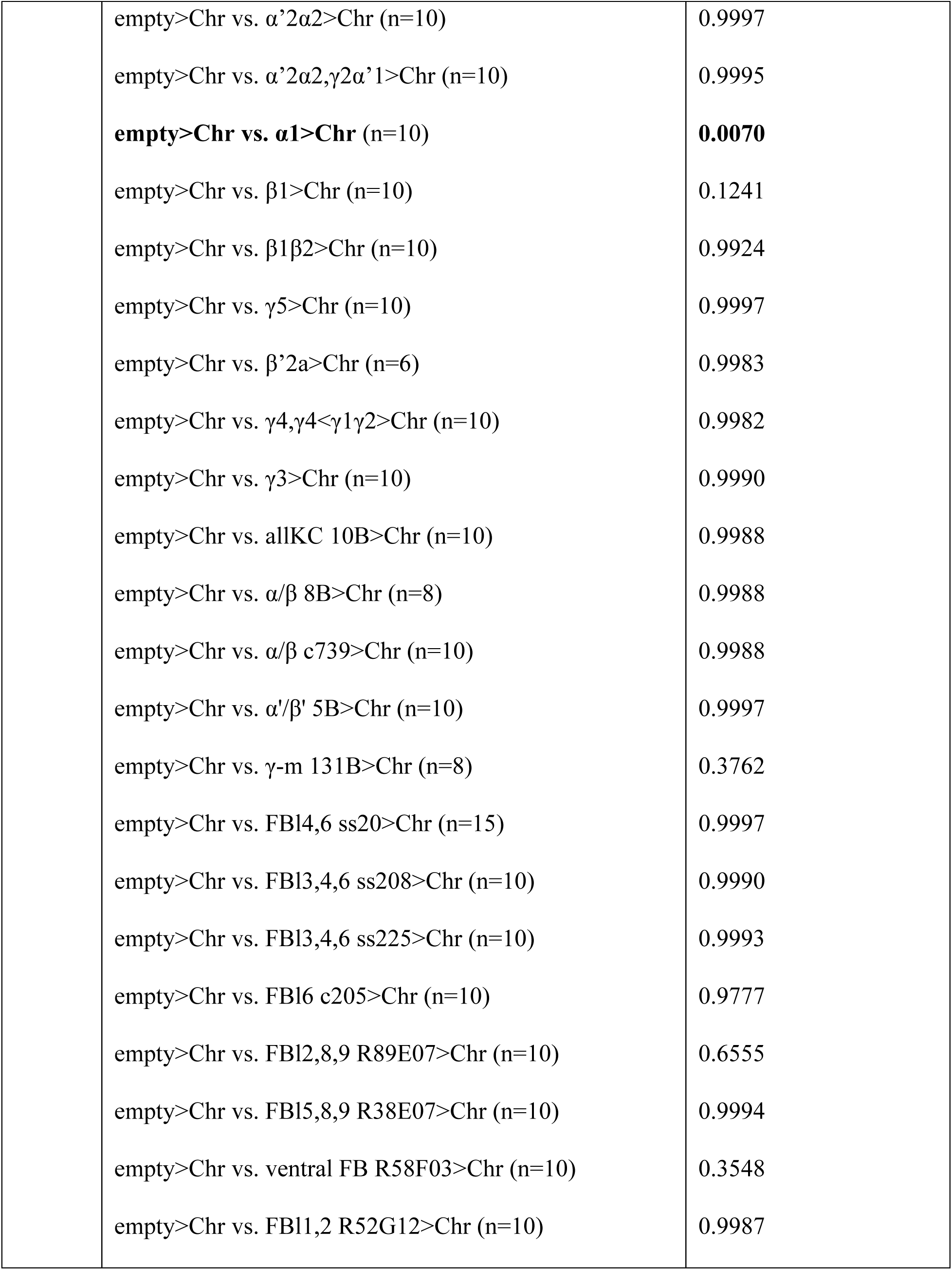
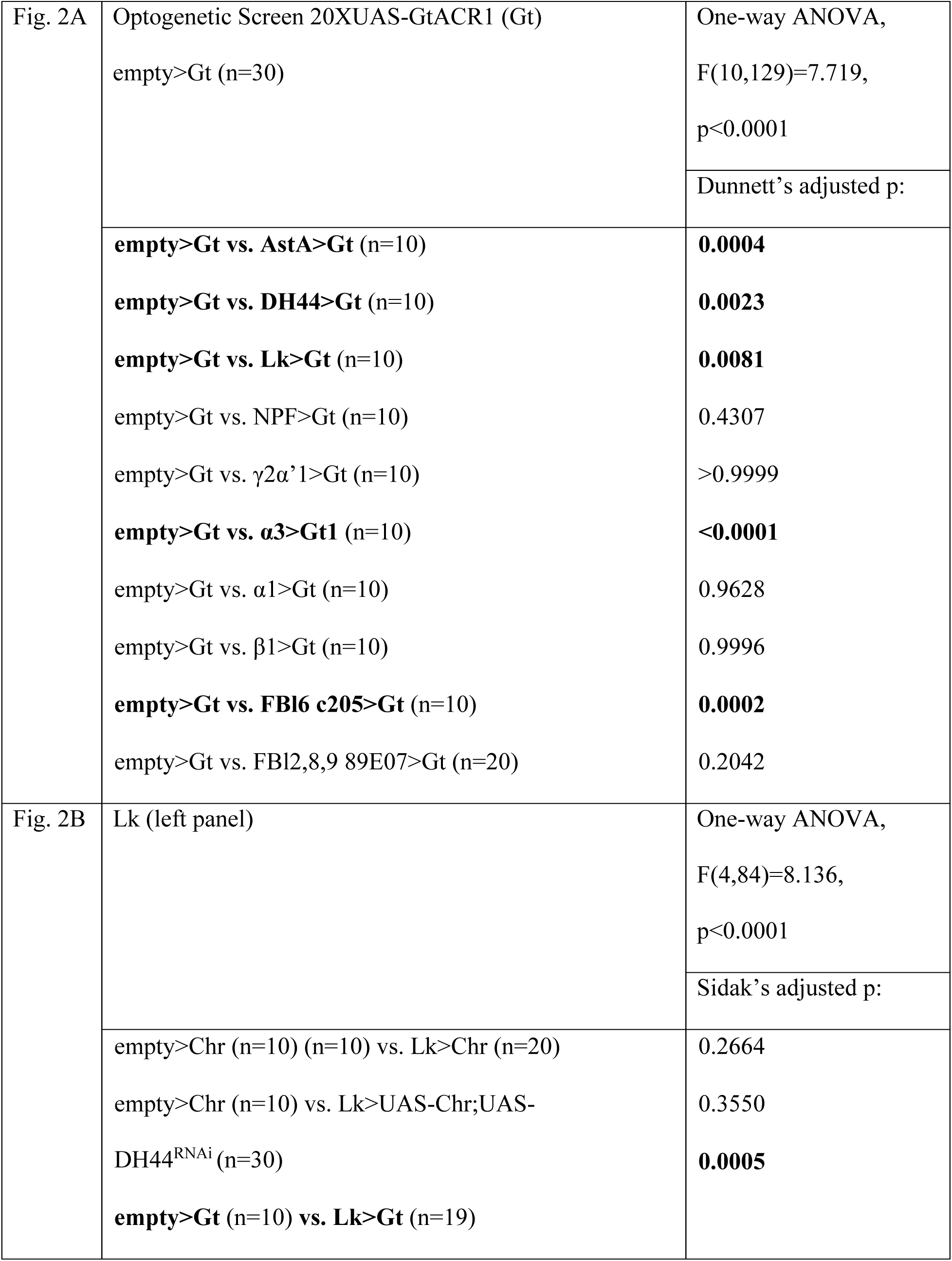
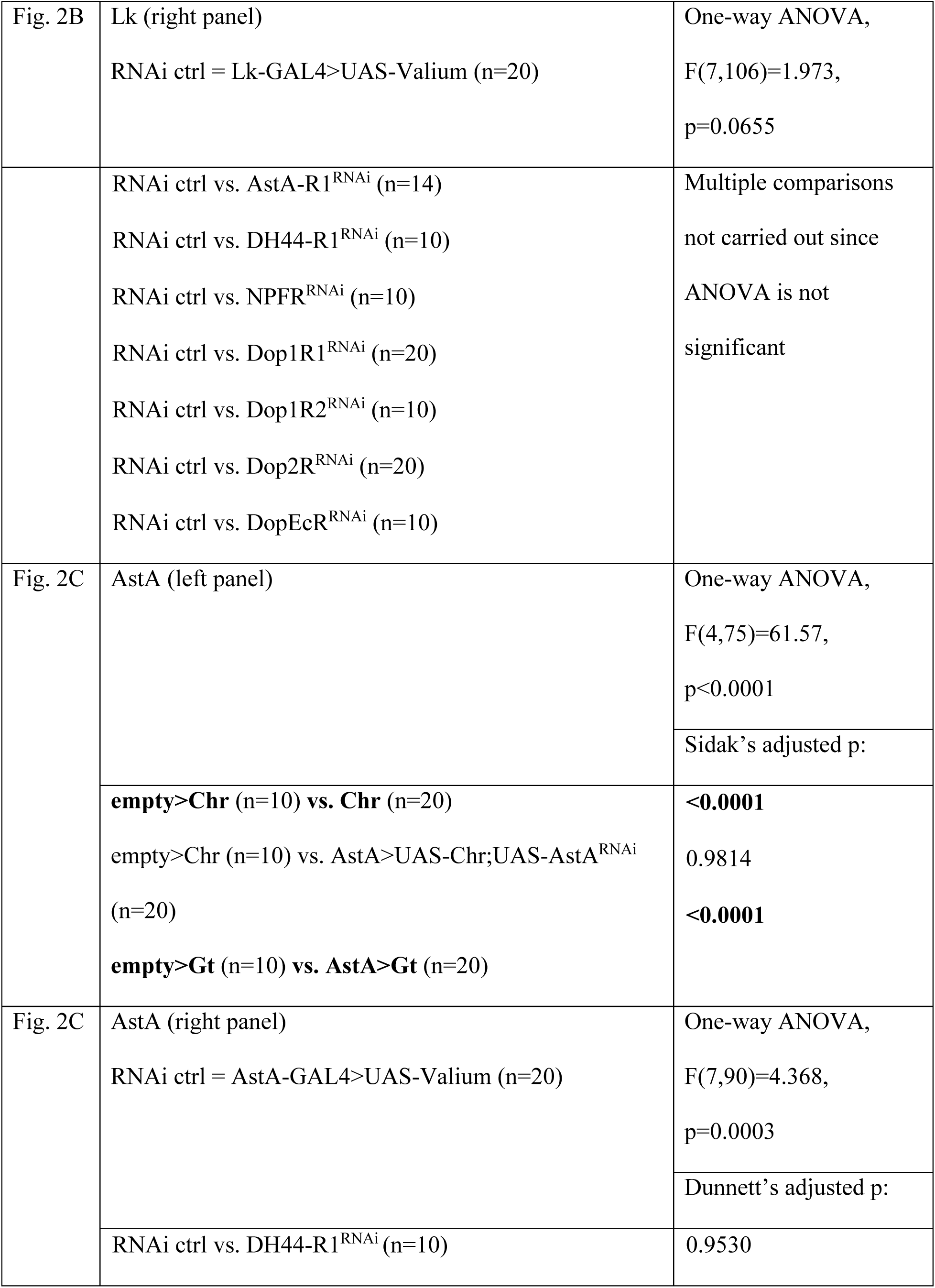
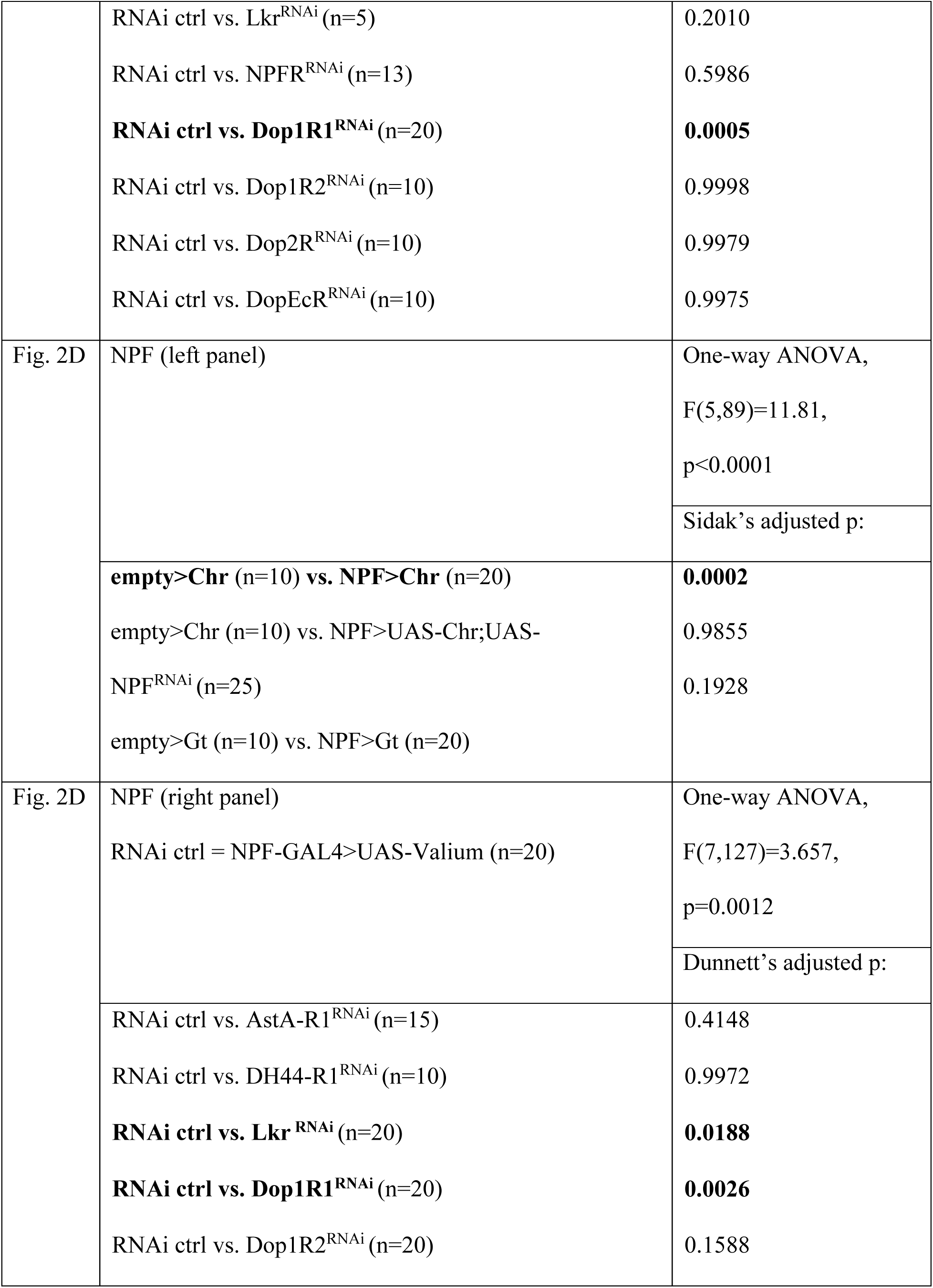
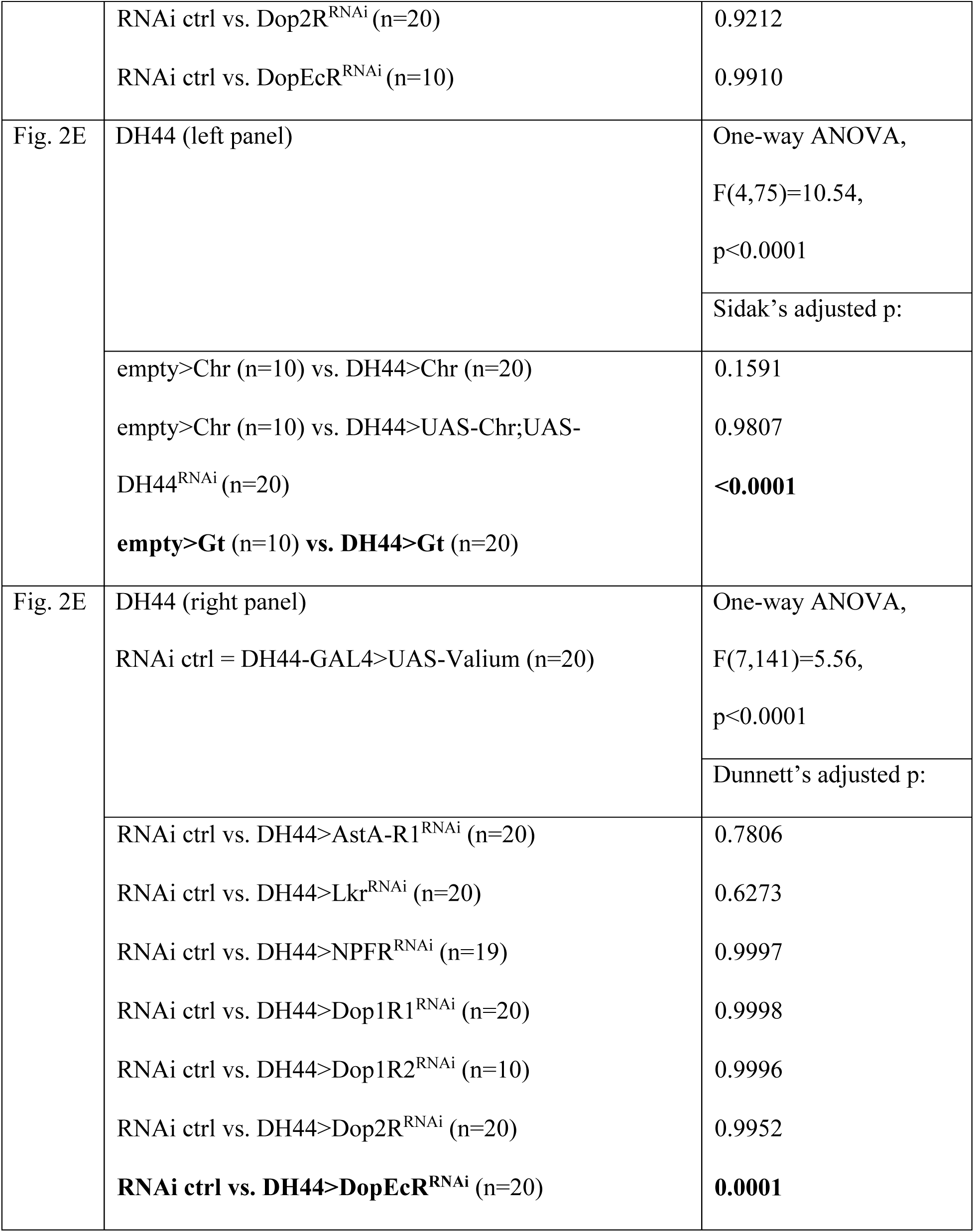
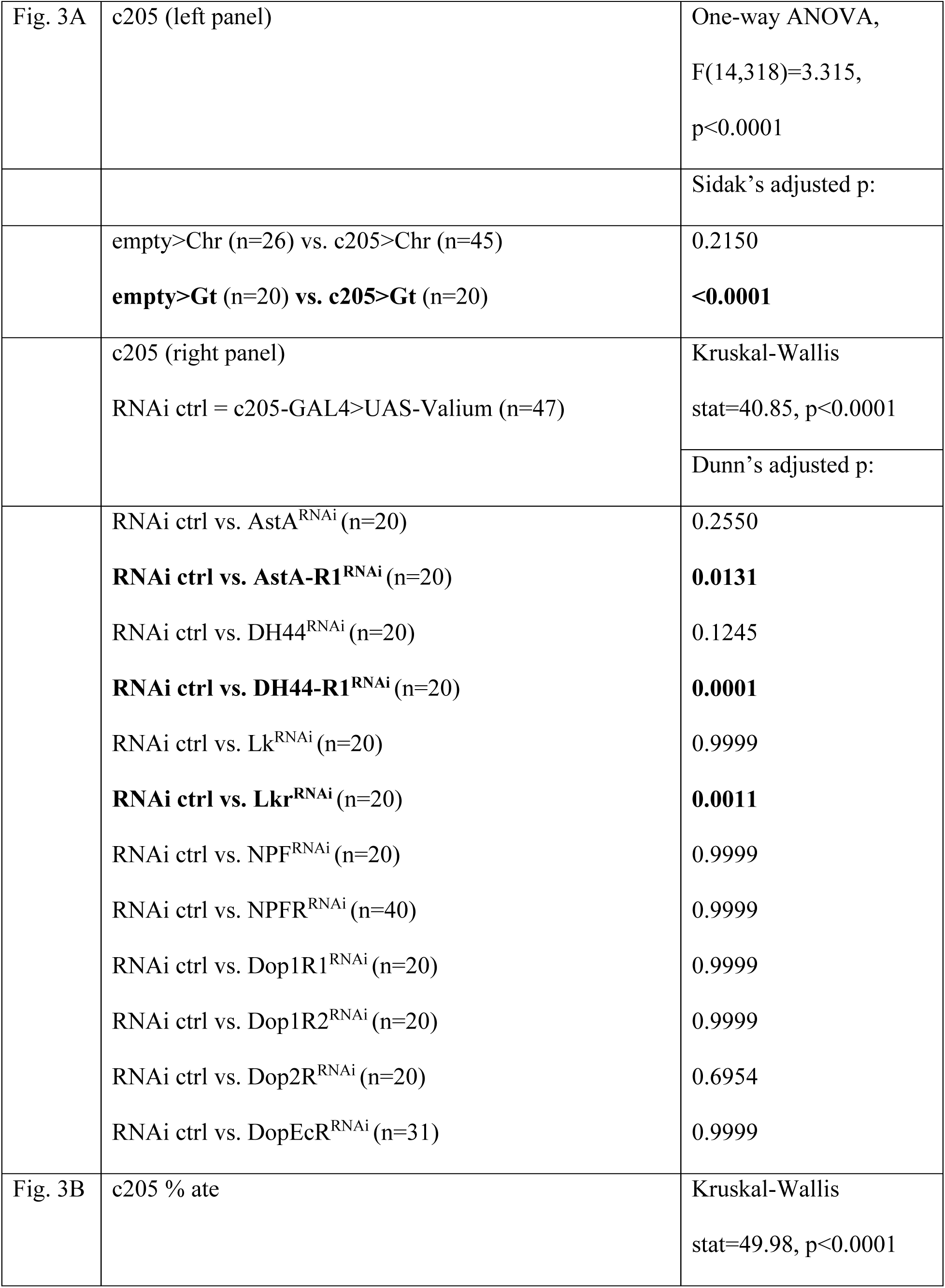
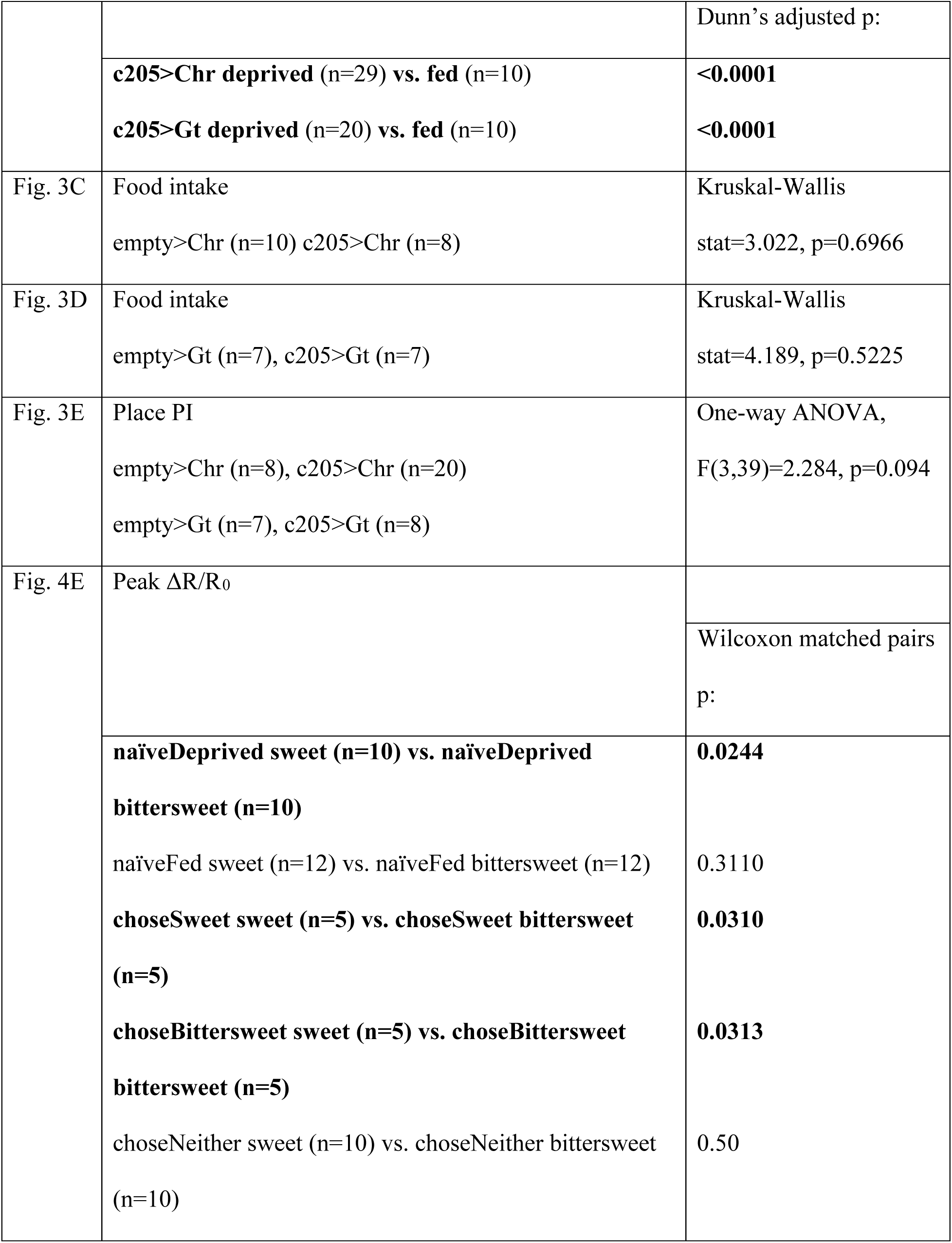
Detailed statistics and sample size for data in the main figures

**Table S2.**
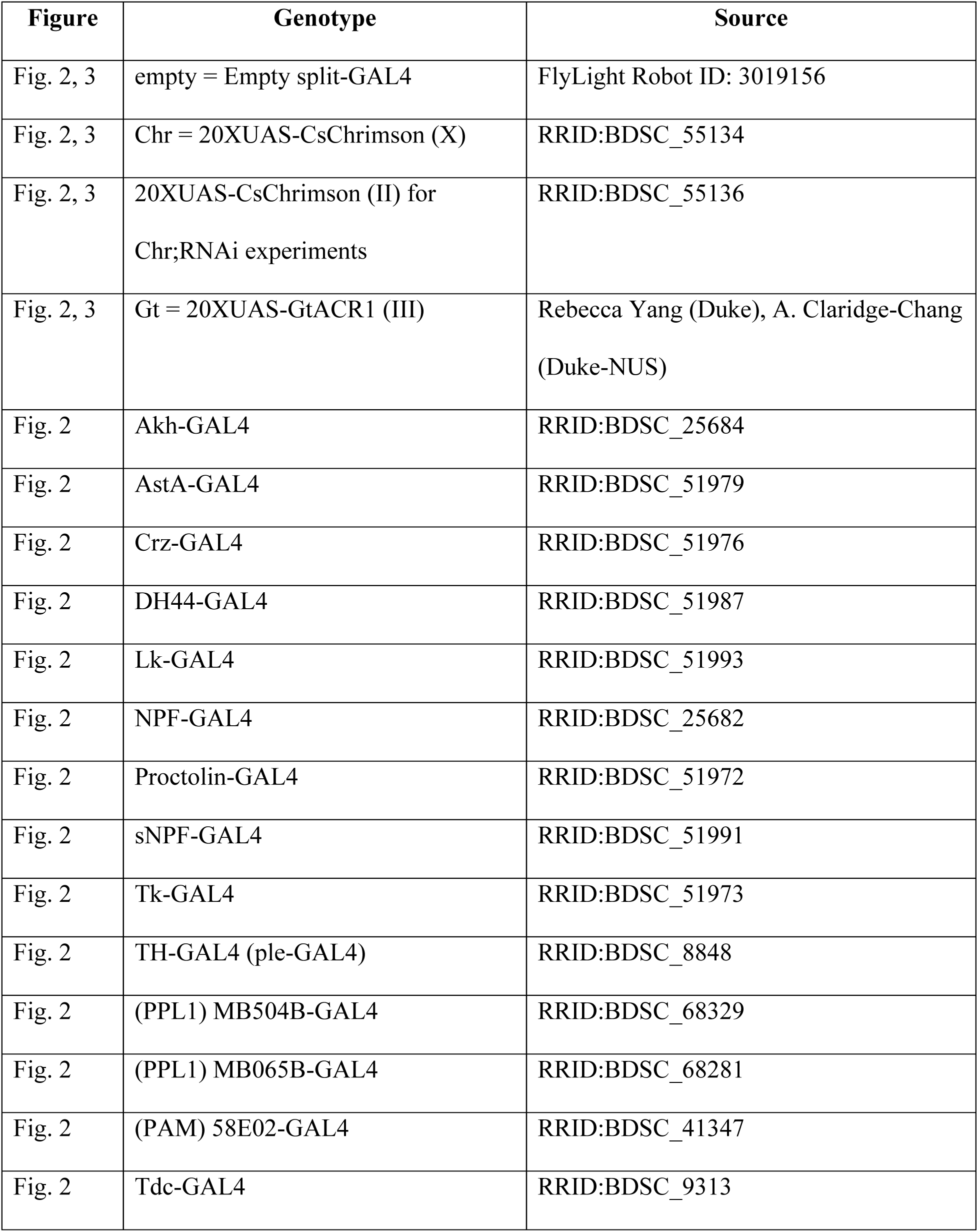
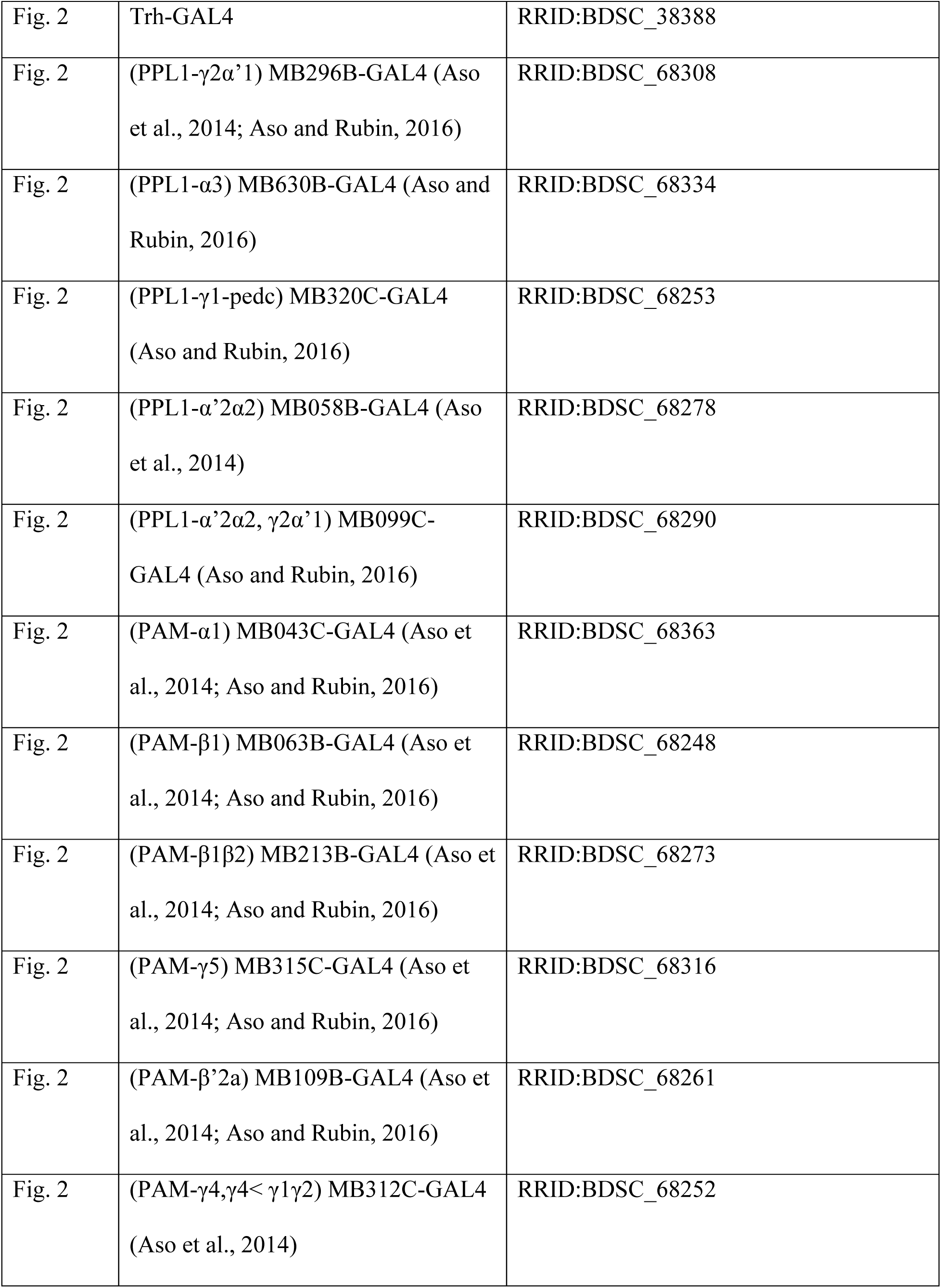
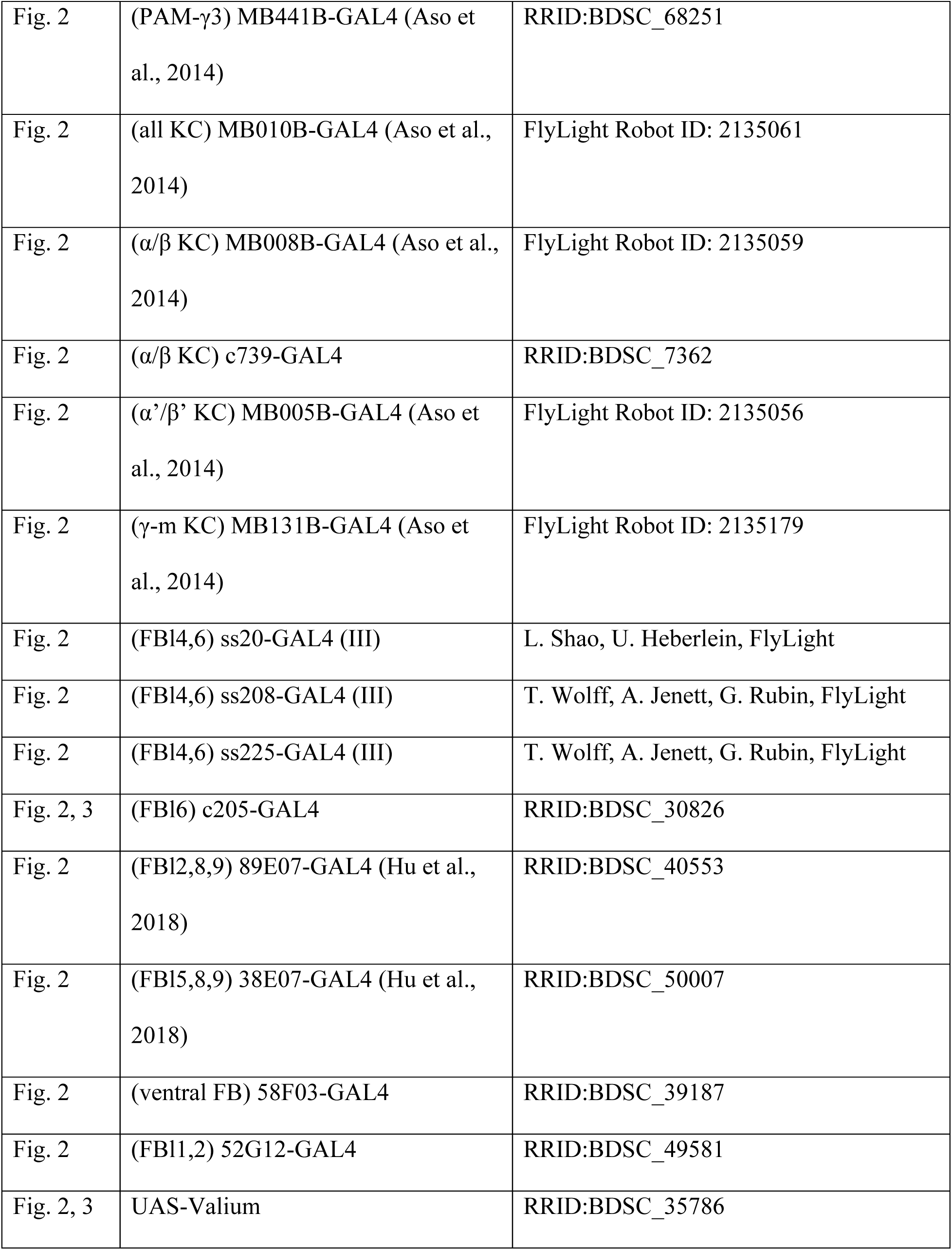
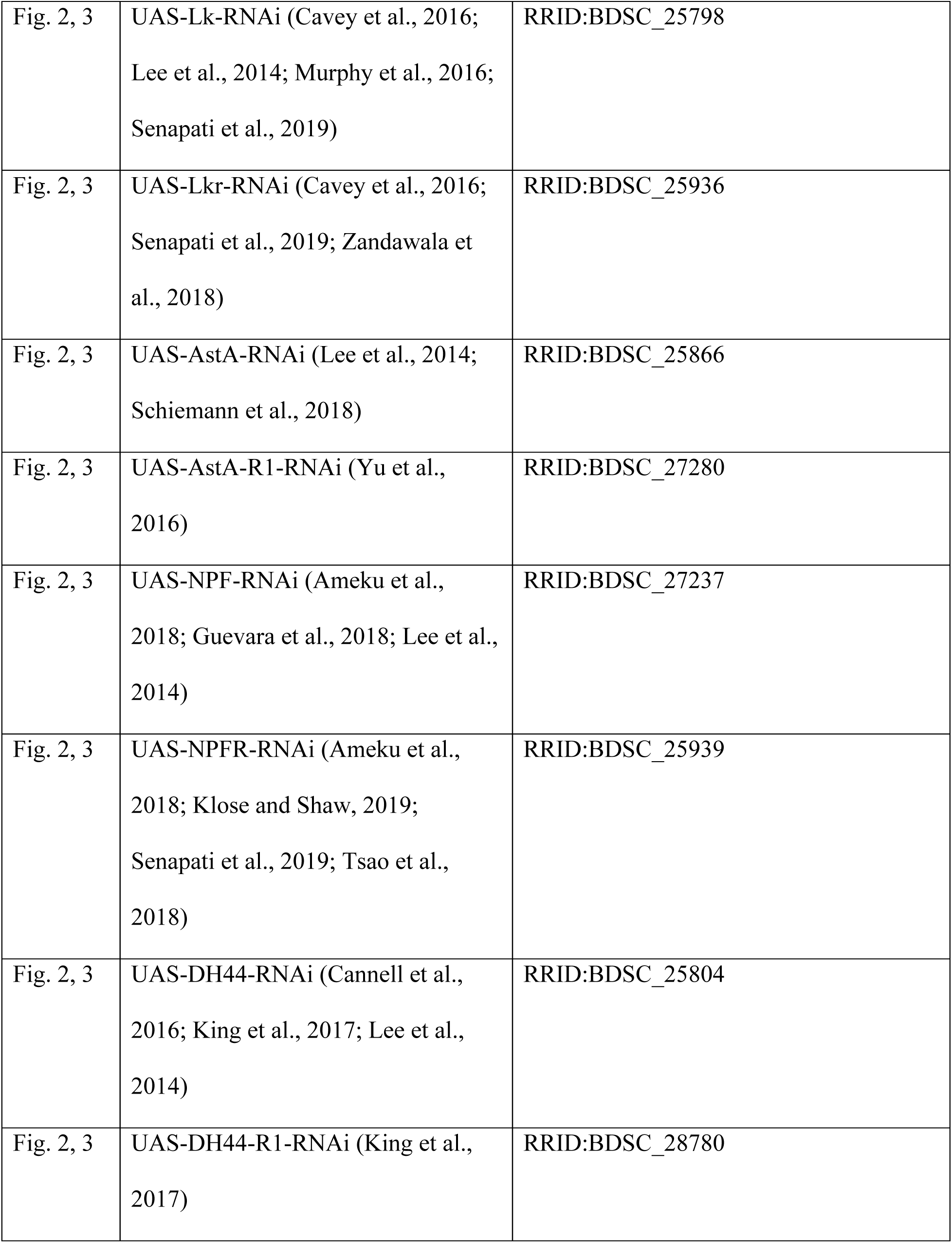
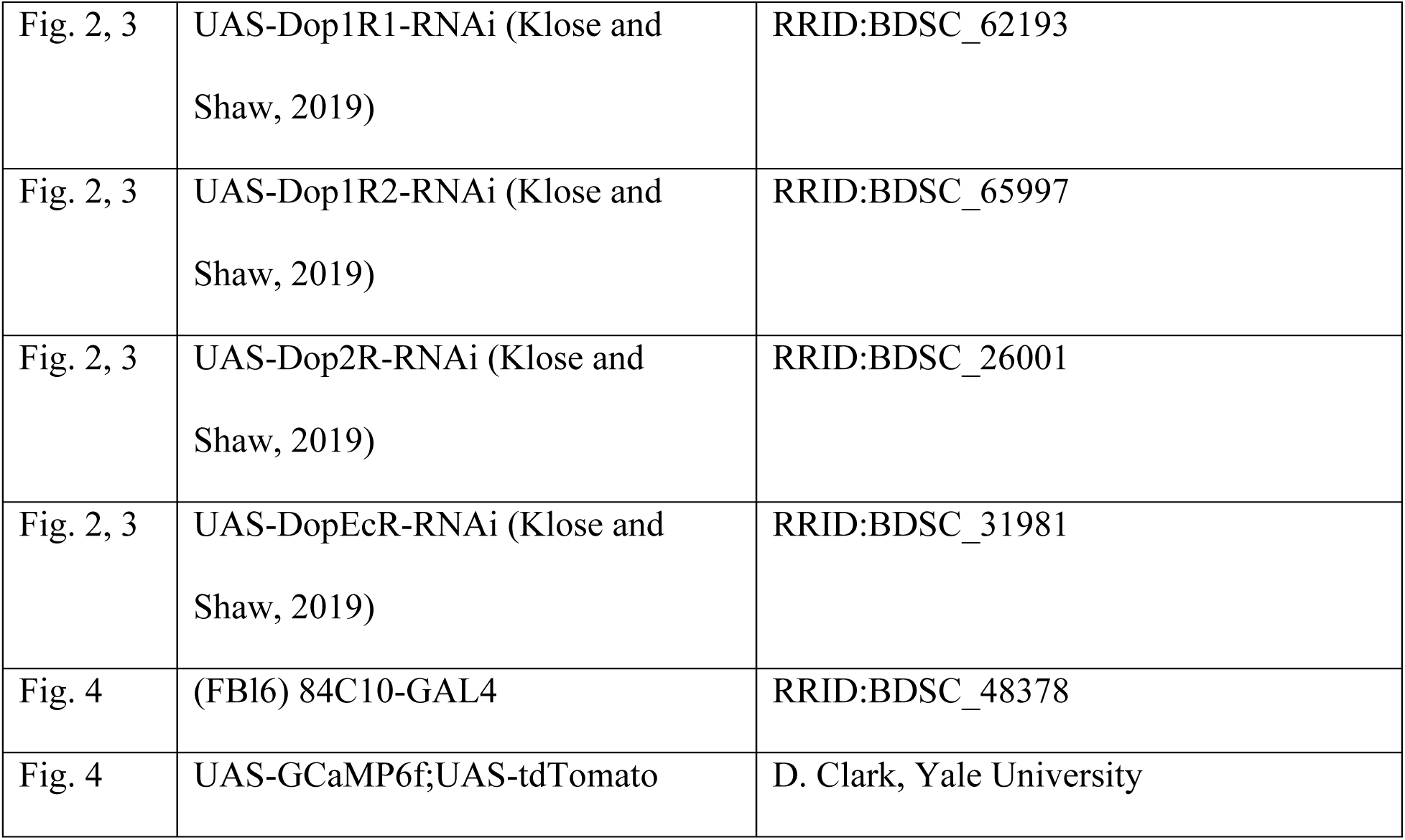
Source for all the fly genotypes used

**Table S3.** Detailed statistics and sample size for data in the supplemental figures

| Figure | Datasets compared | Statistics |
| --- | --- | --- |
| Fig. S3B | 84C10-GAL4 Preference index | One-way ANOVA,<br>F(3,50)=7.823,<br>p=0.0002 |
|  |  | Sidak's adjusted p: |
|  | 84C10>Valium (n=17) vs. 84C10>Chr (n=17)<br>**84C10>Valium (n=10) vs. 84C10>Gt (n=10)** | 0.1463<br>**0.0034** |
| Fig. S3C | 84C10-GAL4 % ate | One-way ANOVA,<br>F(3,55)=186.1,<br>p<0.0001 |
|  |  | Sidak's adjusted p: |
|  | deprived 84C10>Chr (n=17) vs. fed 84C10>Chr<br>(n=15) | <0.0001 |
|  | deprived 84C10>Gt (n=20) vs. fed 84C10>Gt<br>(n=7) | <0.0001 |

## Supplementary Figure Legends

**Figure S1.**
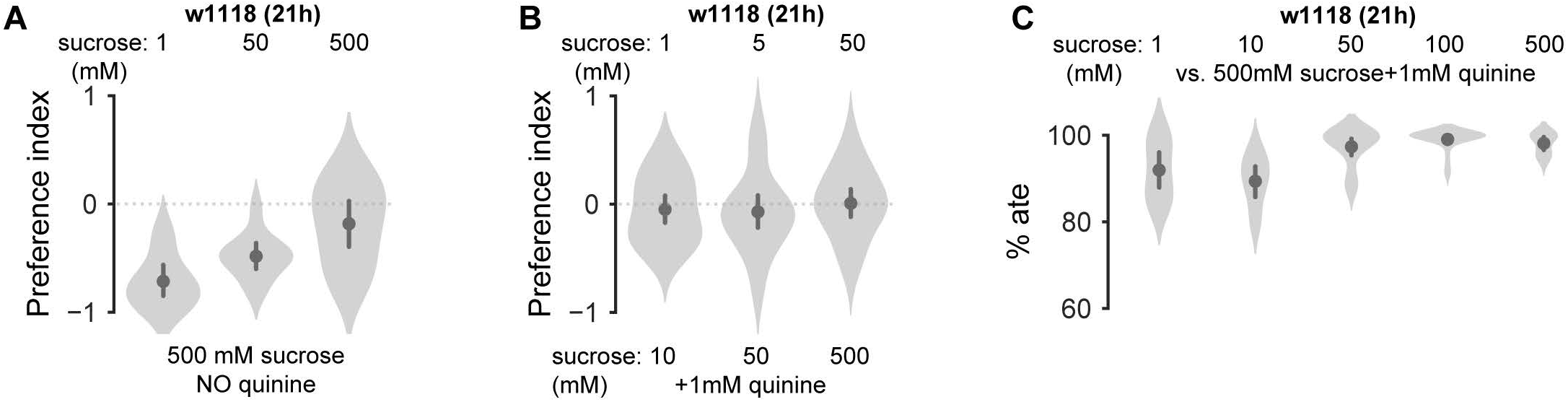
Wild-type fly behavior. **A**, w1118 flies always preferred higher sucrose concentration when no quinine was present. **B**, Food preference was sucrose concentration ratio dependent between two food options when quinine concentration was kept constant in the bittersweet food. **C**, Most w1118 flies ate at 21 h food deprivation, with almost 100% eating at the equal-preference 50 mM sucrose condition. Plots depict mean ± 95% CI; violins show data distribution.

**Figure S2.**
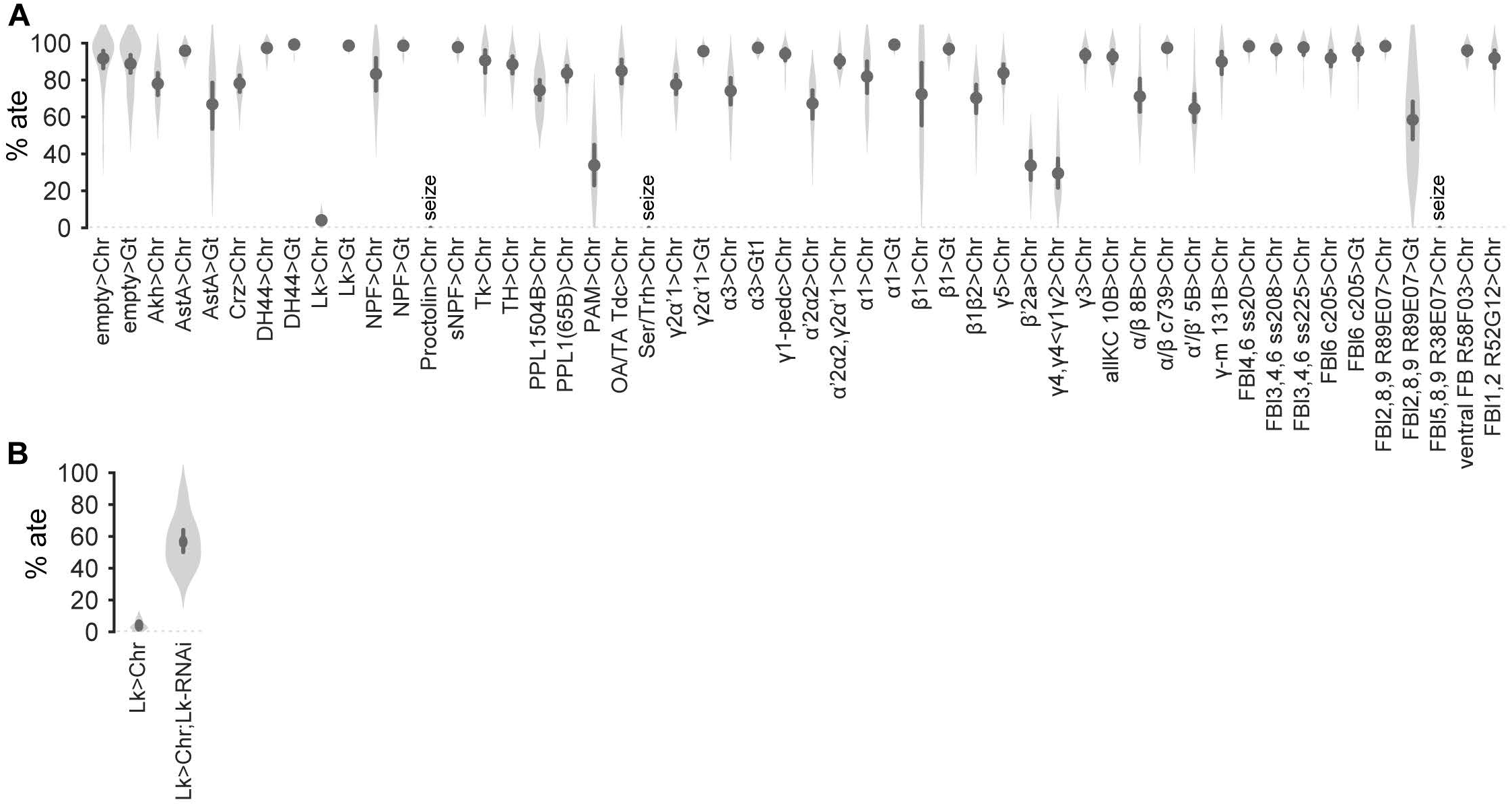
Percentage of flies that ate during decision assay. **A**, Percent of flies that ate during the optogenetic screen for all the genotypes tested. **B**, Only ∼4% of the flies ate when Lk neurons were activated (Lk>Chr) and this effect was abolished (∼57% ate) by knocking down Lk in the same neurons during optogenetic activation (Lk>Chr;Lk-RNAi). Plots depict mean ± 95% CI; violins show data distribution.

**Figure S3.**
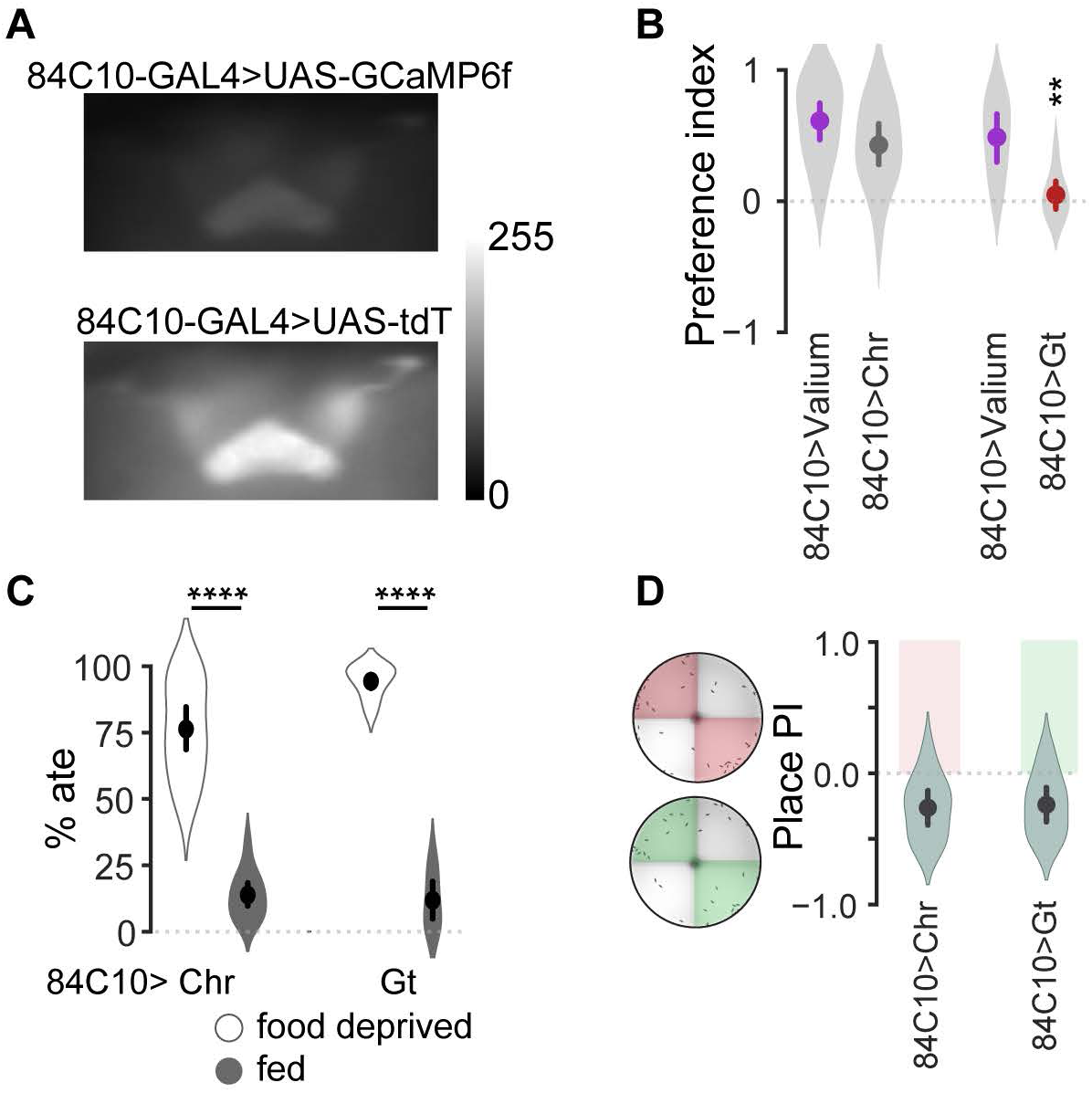
84C10-GAL4 characterization. **A**, 84C10-GAL4 showed high baseline GCaMP6f fluorescence. Images shown are raw florescence images from the same frame without background subtraction. **B**, 84C10-GAL4 had the same behavioral phenotype as c205-GAL4 when optogenetically activated (84C10>Chr) and inhibited (84C10>Gt) compared to controls. Flies prefered bittersweet food compared to control flies when FBl6 neurons were inhibited using 84C10-GAL4. **C**, Feeding was not initiated in fed flies and not inhibited in food-deprived flies on FBl6 activation or inhibition. **D**, Neither activation nor inhibition of FBl6 was inherently rewarding or aversive since there was no significant difference in place preference without food for illuminated vs. non-illuminated sectors of the arena. Plots depict mean ± 95% CI; violins show data distribution. See Table S3 for statistics and sample size. p<0.00001=****, p<0.0001=***, p<0.01=**, p<0.05=*.

**Figure S4.**
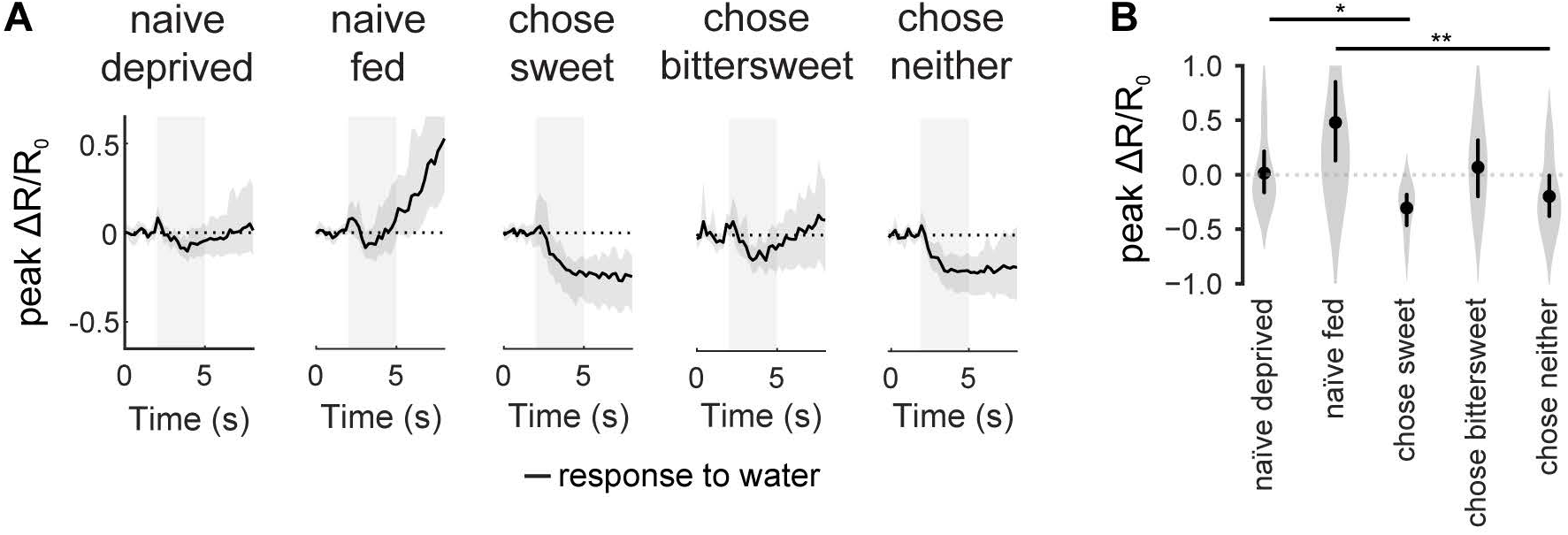
FBl6 neurons respond to water choice. **A**, Ratiometric calcium responses to water stimuli, ΔR/R_0_, of flies with different hunger state and decision outcomes. FBl6 water responses were also modulated by hunger and past experience. FBl6 neurons of food-deprived flies that experienced the decision assay and chose sweet (“chose sweet”) or neither (“chose neither”) food were strongly inhibited by water, while FBl6 neurons of naïve flies were not (“naïve deprived”, “naïve fed”), even though FBl6 taste responses were similar in these flies (Fig. 4d-e). FBl6 neurons of food-deprived flies that chose bittersweet were not inhibited by water (“chose bittersweet”). Gray background area represents water application. Calcium activity trace depicts mean ΔR/R_0_ ± 95% CI. **B**, Peak ΔR/R_0_ shows significant difference between response to rejected vs. chosen water stimuli. p<0.01=**, p<0.05=* (see Table S1 for details on statistics and sample size). Points on graphs depict mean ± 95% CI, with violins depicting full data distribution.

**Figure S5.**
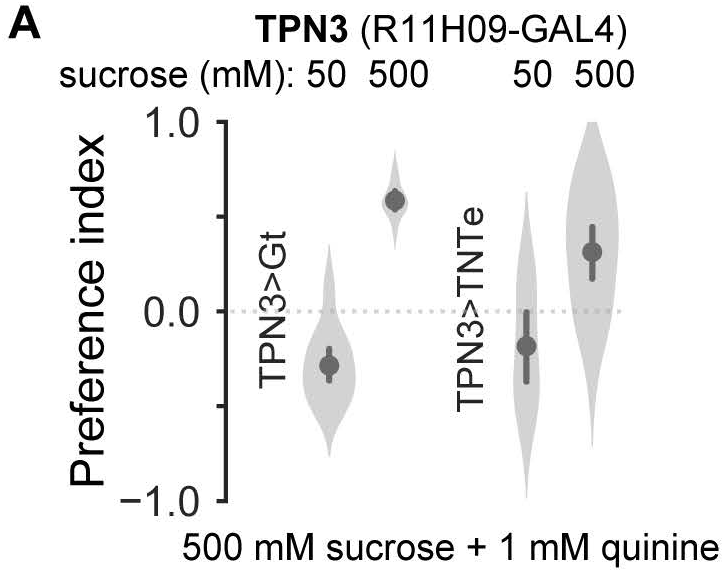
Inhibition of TPN3 bitter taste projection neurons. **A**, Acute optogenetic (TPN3>Gt) and chronic inhibition (TPN3>TNTe) of second-order taste projection neurons (TPN3) did not change food preference at the equal-preference condition (50 mM) or when sucrose concentration is the same in both sweet and bittersweet options (500 mM).

## References

Al-Anzi, B., Armand, E., Nagamei, P., Olszewski, M., Sapin, V., Waters, C., Zinn, K., Wyman, R.J., and Benzer, S. (2010). The leucokinin pathway and its neurons regulate meal size in Drosophila. Curr Biol 20, 969–978.

Aso, Y., Sitaraman, D., Ichinose, T., Kaun, K.R., Vogt, K., Belliart-Guerin, G., Placais, P.Y., Robie, A.A., Yamagata, N., Schnaitmann, C., et al. (2014). Mushroom body output neurons encode valence and guide memory-based action selection in Drosophila. Elife 3, e04580.

Azanchi, R., Kaun, K.R., and Heberlein, U. (2013). Competing dopamine neurons drive oviposition choice for ethanol in Drosophila. Proc Natl Acad Sci U S A 110, 21153–21158.

Berry, J.A., Cervantes-Sandoval, I., Nicholas, E.P., and Davis, R.L. (2012). Dopamine is required for learning and forgetting in Drosophila. Neuron 74, 530–542.

Bohra, A.A., Kallman, B.R., Reichert, H., and VijayRaghavan, K. (2018). Identification of a Single Pair of Interneurons for Bitter Taste Processing in the Drosophila Brain. Curr Biol 28, 847–858 e843.

Brand, A.H., and Perrimon, N. (1993). Targeted gene expression as a means of altering cell fates and generating dominant phenotypes. Development 118, 401–415.

Cannell, E., Dornan, A.J., Halberg, K.A., Terhzaz, S., Dow, J.A.T., and Davies, S.A. (2016). The corticotropin-releasing factor-like diuretic hormone 44 (DH44) and kinin neuropeptides modulate desiccation and starvation tolerance in Drosophila melanogaster. Peptides 80, 96–107.

Cavey, M., Collins, B., Bertet, C., and Blau, J. (2016). Circadian rhythms in neuronal activity propagate through output circuits. Nat Neurosci 19, 587–595.

Chen, T.W., Wardill, T.J., Sun, Y., Pulver, S.R., Renninger, S.L., Baohan, A., Schreiter, E.R., Kerr, R.A., Orger, M.B., Jayaraman, V., et al. (2013). Ultrasensitive fluorescent proteins for imaging neuronal activity. Nature 499, 295–300.

Chu, B., Chui, V., Mann, K., and Gordon, M.D. (2014). Presynaptic gain control drives sweet and bitter taste integration in Drosophila. Curr Biol 24, 1978–1984.

Cohn, R., Morantte, I., and Ruta, V. (2015). Coordinated and Compartmentalized Neuromodulation Shapes Sensory Processing in Drosophila. Cell 163, 1742–1755.

Deng, B., Li, Q., Liu, X., Cao, Y., Li, B., Qian, Y., Xu, R., Mao, R., Zhou, E., Zhang, W., et al. (2019). Chemoconnectomics: Mapping Chemical Transmission in Drosophila. Neuron 101, 876–893 e874.

Deshpande, S.A., Carvalho, G.B., Amador, A., Phillips, A.M., Hoxha, S., Lizotte, K.J., and Ja, W.W. (2014). Quantifying Drosophila food intake: comparative analysis of current methodology. Nat Methods 11, 535–540.

Donlea, J.M., Pimentel, D., and Miesenbock, G. (2014). Neuronal machinery of sleep homeostasis in Drosophila. Neuron 81, 860–872.

Donlea, J.M., Pimentel, D., Talbot, C.B., Kempf, A., Omoto, J.J., Hartenstein, V., and Miesenbock, G. (2018). Recurrent Circuitry for Balancing Sleep Need and Sleep. Neuron 97, 378–389 e374.

Dus, M., Lai, J.S., Gunapala, K.M., Min, S., Tayler, T.D., Hergarden, A.C., Geraud, E., Joseph, C.M., and Suh, G.S. (2015). Nutrient Sensor in the Brain Directs the Action of the Brain-Gut Axis in Drosophila. Neuron 87, 139–151.

FlyLight, J. Janelia FlyLight. Giraldo, Y.M., Leitch, K.J., Ros, I.G., Warren, T.L., Weir, P.T., and Dickinson, M.H. (2018). Sun Navigation Requires Compass Neurons in Drosophila. Curr Biol 28, 2845–2852 e2844.

Gold, J.I., and Shadlen, M.N. (2007). The neural basis of decision making. Annu Rev Neurosci 30, 535–574.

Green, J., Vijayan, V., Mussells Pires, P., Adachi, A., and Maimon, G. (2019). A neural heading estimate is compared with an internal goal to guide oriented navigation. Nat Neurosci 22, 1460–1468.

Guo, F., Holla, M., Diaz, M.M., and Rosbash, M. (2018). A Circadian Output Circuit Controls Sleep-Wake Arousal in Drosophila. Neuron 100, 624–635 e624.

Hentze, J.L., Carlsson, M.A., Kondo, S., Nassel, D.R., and Rewitz, K.F. (2015). The Neuropeptide Allatostatin A Regulates Metabolism and Feeding Decisions in Drosophila. Sci Rep 5, 11680.

Honkanen, A., Adden, A., da Silva Freitas, J., and Heinze, S. (2019). The insect central complex and the neural basis of navigational strategies. J Exp Biol 222.

Huetteroth, W., Perisse, E., Lin, S., Klappenbach, M., Burke, C., and Waddell, S. (2015). Sweet taste and nutrient value subdivide rewarding dopaminergic neurons in Drosophila. Curr Biol 25, 751–758.

Jenett, A., Rubin, G.M., Ngo, T.T., Shepherd, D., Murphy, C., Dionne, H., Pfeiffer, B.D., Cavallaro, A., Hall, D., Jeter, J., et al. (2012). A GAL4-driver line resource for Drosophila neurobiology. Cell Rep 2, 991–1001.

Jeong, Y.T., Shim, J., Oh, S.R., Yoon, H.I., Kim, C.H., Moon, S.J., and Montell, C. (2013). An odorant-binding protein required for suppression of sweet taste by bitter chemicals. Neuron 79, 725–737.

Jody, C., Cristian, G., Antje, K., Hideo, O., Rob, S., and Konrad, R. (2020). NeuronBridge Codebase.

Kahsai, L., and Winther, A.M. (2011). Chemical neuroanatomy of the Drosophila central complex: distribution of multiple neuropeptides in relation to neurotransmitters. J Comp Neurol 519, 290–315.

Kain, P., and Dahanukar, A. (2015). Secondary taste neurons that convey sweet taste and starvation in the Drosophila brain. Neuron 85, 819–832.

Kim, H., Kirkhart, C., and Scott, K. (2017a). Long-range projection neurons in the taste circuit of Drosophila. Elife 6.

Kim, J.H., Ki, Y., Lee, H., Hur, M.S., Baik, B., Hur, J.H., Nam, D., and Lim, C. (2020). The voltage-gated potassium channel Shaker promotes sleep via thermosensitive GABA transmission. Commun Biol 3, 174.

Kim, S.S., Rouault, H., Druckmann, S., and Jayaraman, V. (2017b). Ring attractor dynamics in the Drosophila central brain. Science 356, 849–853.

Kirkhart, C., and Scott, K. (2015). Gustatory learning and processing in the Drosophila mushroom bodies. J Neurosci 35, 5950–5958.

Klapoetke, N.C., Murata, Y., Kim, S.S., Pulver, S.R., Birdsey-Benson, A., Cho, Y.K., Morimoto, T.K., Chuong, A.S., Carpenter, E.J., Tian, Z., et al. (2014). Independent optical excitation of distinct neural populations. Nat Methods 11, 338–346.

Lee, D., Seo, H., and Jung, M.W. (2012). Neural basis of reinforcement learning and decision making. Annu Rev Neurosci 35, 287–308.

Lewis, L.P., Siju, K.P., Aso, Y., Friedrich, A.B., Bulteel, A.J., Rubin, G.M., and Grunwald Kadow, I.C. (2015). A Higher Brain Circuit for Immediate Integration of Conflicting Sensory Information in Drosophila. Curr Biol 25, 2203–2214.

Lin, S., Senapati, B., and Tsao, C.H. (2019). Neural basis of hunger-driven behaviour in Drosophila. Open Biol 9, 180259.

Liu, C., Meng, Z., Wiggin, T.D., Yu, J., Reed, M.L., Guo, F., Zhang, Y., Rosbash, M., and Griffith, L.C. (2019). A Serotonin-Modulated Circuit Controls Sleep Architecture to Regulate Cognitive Function Independent of Total Sleep in Drosophila. Curr Biol 29, 3635–3646 e3635.

Liu, C., Placais, P.Y., Yamagata, N., Pfeiffer, B.D., Aso, Y., Friedrich, A.B., Siwanowicz, I., Rubin, G.M., Preat, T., and Tanimoto, H. (2012a). A subset of dopamine neurons signals reward for odour memory in Drosophila. Nature 488, 512–516.

Liu, Q., Liu, S., Kodama, L., Driscoll, M.R., and Wu, M.N. (2012b). Two dopaminergic neurons signal to the dorsal fan-shaped body to promote wakefulness in Drosophila. Curr Biol 22, 2114–2123.

Liu, S., Liu, Q., Tabuchi, M., and Wu, M.N. (2016). Sleep Drive Is Encoded by Neural Plastic Changes in a Dedicated Circuit. Cell 165, 1347–1360.

Lyutova, R., Selcho, M., Pfeuffer, M., Segebarth, D., Habenstein, J., Rohwedder, A., Frantzmann, F., Wegener, C., Thum, A.S., and Pauls, D. (2019). Reward signaling in a recurrent circuit of dopaminergic neurons and peptidergic Kenyon cells. Nat Commun 10, 3097.

Masek, P., and Scott, K. (2010). Limited taste discrimination in Drosophila. Proc Natl Acad Sci U S A 107, 14833–14838.

Masek, P., Worden, K., Aso, Y., Rubin, G.M., and Keene, A.C. (2015). A dopamine-modulated neural circuit regulating aversive taste memory in Drosophila. Curr Biol 25, 1535–1541.

Mathejczyk, T.F., and Wernet, M.F. (2019). Heading choices of flying Drosophila under changing angles of polarized light. Sci Rep 9, 16773.

Meissner, G.W., Dorman, Z., Nern, A., Forster, K., Gibney, T., Jeter, J., Johnson, L., He, Y., Lee, K., Melton, B., et al. (2020). An image resource of subdivided <em>Drosophila</em> GAL4-driver expression patterns for neuron-level searches. bioRxiv, 2020.2005.2029.080473.

Meunier, N., Marion-Poll, F., Rospars, J.P., and Tanimura, T. (2003). Peripheral coding of bitter taste in Drosophila. J Neurobiol 56, 139–152.

Miyazaki, T., Lin, T.Y., Ito, K., Lee, C.H., and Stopfer, M. (2015). A gustatory second-order neuron that connects sucrose-sensitive primary neurons and a distinct region of the gnathal ganglion in the Drosophila brain. J Neurogenet 29, 144–155.

Mohammad, F., Stewart, J.C., Ott, S., Chlebikova, K., Chua, J.Y., Koh, T.W., Ho, J., and Claridge-Chang, A. (2017). Optogenetic inhibition of behavior with anion channelrhodopsins. Nat Methods 14, 271–274.

Nassel, D.R., and Zandawala, M. (2019). Recent advances in neuropeptide signaling in Drosophila, from genes to physiology and behavior. Prog Neurobiol 179, 101607.

Ni, J.Q., Liu, L.P., Binari, R., Hardy, R., Shim, H.S., Cavallaro, A., Booker, M., Pfeiffer, B.D., Markstein, M., Wang, H., et al. (2009). A Drosophila resource of transgenic RNAi lines for neurogenetics. Genetics 182, 1089–1100.

Park, I.M., Meister, M.L., Huk, A.C., and Pillow, J.W. (2014). Encoding and decoding in parietal cortex during sensorimotor decision-making. Nat Neurosci 17, 1395–1403.

Perkins, L.A., Holderbaum, L., Tao, R., Hu, Y., Sopko, R., McCall, K., Yang-Zhou, D., Flockhart, I., Binari, R., Shim, H.-S., et al. (2015). The Transgenic RNAi Project at Harvard Medical School: Resources and Validation. Genetics 201, 843–852.

Pimentel, D., Donlea, J.M., Talbot, C.B., Song, S.M., Thurston, A.J.F., and Miesenbock, G. (2016). Operation of a homeostatic sleep switch. Nature 536, 333–337.

Qian, Y., Cao, Y., Deng, B., Yang, G., Li, J., Xu, R., Zhang, D., Huang, J., and Rao, Y. (2017). Sleep homeostasis regulated by 5HT2b receptor in a small subset of neurons in the dorsal fan-shaped body of drosophila. Elife 6.

Ramdya, P., Schneider, J., and Levine, J.D. (2017). The neurogenetics of group behavior in Drosophila melanogaster. J Exp Biol 220, 35–41.

Riemensperger, T., Voller, T., Stock, P., Buchner, E., and Fiala, A. (2005). Punishment prediction by dopaminergic neurons in Drosophila. Curr Biol 15, 1953–1960.

Rooke, R., Rasool, A., Schneider, J., and Levine, J.D. (2020). Drosophila melanogaster behaviour changes in different social environments based on group size and density. Commun Biol 3, 304.

Scaplen, K.M., Talay, M., Nunez, K.M., Salamon, S., Waterman, A.G., Gang, S., Song, S.L., Barnea, G., and Kaun, K.R. (2020). Circuits that encode and guide alcohol-associated preference. Elife 9.

Scott, K. (2018). Gustatory Processing in Drosophila melanogaster. Annu Rev Entomol 63, 15–30.

Seelig, J.D., and Jayaraman, V. (2015). Neural dynamics for landmark orientation and angular path integration. Nature 521, 186–191.

Shadlen, M.N., and Kiani, R. (2013). Decision making as a window on cognition. Neuron 80, 791–806.

Shao, L., Saver, M., Chung, P., Ren, Q., Lee, T., Kent, C.F., and Heberlein, U. (2017). Dissection of the Drosophila neuropeptide F circuit using a high-throughput two-choice assay. Proc Natl Acad Sci U S A 114, E8091–E8099.

Sugrue, L.P., Corrado, G.S., and Newsome, W.T. (2005). Choosing the greater of two goods: neural currencies for valuation and decision making. Nat Rev Neurosci 6, 363–375.

Sun, Y., Nern, A., Franconville, R., Dana, H., Schreiter, E.R., Looger, L.L., Svoboda, K., Kim, D.S., Hermundstad, A.M., and Jayaraman, V. (2017). Neural signatures of dynamic stimulus selection in Drosophila. Nat Neurosci 20, 1104–1113.

Talay, M., Richman, E.B., Snell, N.J., Hartmann, G.G., Fisher, J.D., Sorkac, A., Santoyo, J.F., Chou-Freed, C., Nair, N., Johnson, M., et al. (2017). Transsynaptic Mapping of Second-Order Taste Neurons in Flies by trans-Tango. Neuron 96, 783–795 e784.

Tsao, C.H., Chen, C.C., Lin, C.H., Yang, H.Y., and Lin, S. (2018). Drosophila mushroom bodies integrate hunger and satiety signals to control innate food-seeking behavior. Elife 7.

Tsetsos, K., Chater, N., and Usher, M. (2012). Salience driven value integration explains decision biases and preference reversal. Proc Natl Acad Sci U S A 109, 9659–9664.

Turner-Evans, D., Wegener, S., Rouault, H., Franconville, R., Wolff, T., Seelig, J.D., Druckmann, S., and Jayaraman, V. (2017). Angular velocity integration in a fly heading circuit. Elife 6.

Wang, X.J. (2008). Decision making in recurrent neuronal circuits. Neuron 60, 215–234.

Wong, R., Piper, M.D., Wertheim, B., and Partridge, L. (2009). Quantification of food intake in Drosophila. PLoS One 4, e6063.

Wu, Q., Wen, T., Lee, G., Park, J.H., Cai, H.N., and Shen, P. (2003). Developmental control of foraging and social behavior by the Drosophila neuropeptide Y-like system. Neuron 39, 147–161.

Xu, C.S., Januszewski, M., Lu, Z., Takemura, S.-y., Hayworth, K.J., Huang, G., Shinomiya, K., Maitin-Shepard, J., Ackerman, D., Berg, S., et al. (2020). A Connectome of the Adult <em>Drosophila</em> Central Brain. bioRxiv, 2020.2001.2021.911859.

Yamagata, N., Ichinose, T., Aso, Y., Placais, P.Y., Friedrich, A.B., Sima, R.J., Preat, T., Rubin, G.M., and Tanimoto, H. (2015). Distinct dopamine neurons mediate reward signals for short- and long-term memories. Proc Natl Acad Sci U S A 112, 578–583.

Yang, Z., Huang, R., Fu, X., Wang, G., Qi, W., Mao, D., Shi, Z., Shen, W.L., and Wang, L. (2018). A post-ingestive amino acid sensor promotes food consumption in Drosophila. Cell Res 28, 1013–1025.

Zandawala, M., Yurgel, M.E., Texada, M.J., Liao, S., Rewitz, K.F., Keene, A.C., and Nassel, D.R. (2018). Modulation of Drosophila post-feeding physiology and behavior by the neuropeptide leucokinin. PLoS Genet 14, e1007767.

## Supplementary References

Ameku, T., Yoshinari, Y., Texada, M.J., Kondo, S., Amezawa, K., Yoshizaki, G., Shimada-Niwa, Y., and Niwa, R. (2018). Midgut-derived neuropeptide F controls germline stem cell proliferation in a mating-dependent manner. PLoS Biol 16, e2005004.

Aso, Y., Hattori, D., Yu, Y., Johnston, R.M., Iyer, N.A., Ngo, T.T., Dionne, H., Abbott, L.F., Axel, R., Tanimoto, H., et al. (2014). The neuronal architecture of the mushroom body provides a logic for associative learning. Elife 3, e04577.

Aso, Y., and Rubin, G.M. (2016). Dopaminergic neurons write and update memories with cell-type-specific rules. Elife 5.

Guevara, A., Gates, H., Urbina, B., and French, R. (2018). Developmental Ethanol Exposure Causes Reduced Feeding and Reveals a Critical Role for Neuropeptide F in Survival. Front Physiol 9, 237.

Hu, W., Peng, Y., Sun, J., Zhang, F., Zhang, X., Wang, L., Li, Q., and Zhong, Y. (2018). Fan-Shaped Body Neurons in the Drosophila Brain Regulate Both Innate and Conditioned Nociceptive Avoidance. Cell Rep 24, 1573–1584.

King, A.N., Barber, A.F., Smith, A.E., Dreyer, A.P., Sitaraman, D., Nitabach, M.N., Cavanaugh, D.J., and Sehgal, A. (2017). A Peptidergic Circuit Links the Circadian Clock to Locomotor Activity. Curr Biol 27, 1915–1927 e1915.

Klose, M., and Shaw, P. (2019). Sleep-drive reprograms clock neuronal identity through CREB-binding protein induced PDFR expression. bioRxiv, 2019.2012.2012.874628.

Lee, J.H., Bassel-Duby, R., and Olson, E.N. (2014). Heart- and muscle-derived signaling system dependent on MED13 and Wingless controls obesity in Drosophila. Proc Natl Acad Sci U S A 111, 9491–9496.

Murphy, K.R., Deshpande, S.A., Yurgel, M.E., Quinn, J.P., Weissbach, J.L., Keene, A.C., Dawson-Scully, K., Huber, R., Tomchik, S.M., and Ja, W.W. (2016). Postprandial sleep mechanics in Drosophila. Elife 5.

Schiemann, R., Lammers, K., Janz, M., Lohmann, J., Paululat, A., and Meyer, H. (2018). Identification and In Vivo Characterisation of Cardioactive Peptides in Drosophila melanogaster. Int J Mol Sci 20.

Senapati, B., Tsao, C.H., Juan, Y.A., Chiu, T.H., Wu, C.L., Waddell, S., and Lin, S. (2019). A neural mechanism for deprivation state-specific expression of relevant memories in Drosophila. Nat Neurosci 22, 2029–2039.

Yu, Y., Huang, R., Ye, J., Zhang, V., Wu, C., Cheng, G., Jia, J., and Wang, L. (2016). Regulation of starvation-induced hyperactivity by insulin and glucagon signaling in adult Drosophila. Elife 5.

